# Multiple modes of regulation control dynamic transcription patterns during the mitosis-G1 transition

**DOI:** 10.1101/2021.06.22.449286

**Authors:** Luke A. Wojenski, Lauren M. Wainman, Geno Villafano, Chris Kuhlberg, Pariksheet Nanda, Leighton Core

**Author notes:** Current address: The Jackson Laboratory for Genomic Medicine, Farmington, CT, USA 06032.

## Abstract

Following cell division, genomes must reactivate gene expression patterns that reflect the identity of the cell. Here, we use PRO-seq to examine the mechanisms that reestablish transcription patterns after mitosis. We uncover regulation of the transcription cycle at multiple steps including initiation, promoter-proximal pause positioning and escape, poly-A site cleavage and termination during the mitotic-G1 transition. During mitosis, RNA polymerase activity is retained at initiation sites, albeit shifted in position relative to non-mitotic cells. This activity is strongly linked to maintenance of local chromatin architecture during mitosis and is more predictive of rapid gene reactivation than histone modifications previously associated with bookmarking. These molecular bookmarks, combined with sequence-specific transcription factors, direct expression of select cell growth and cell specific genes during mitosis followed by reactivation of functional gene groups with distinct kinetics after mitosis. This study details how dynamic regulation of transcription at multiple steps contributes to gene expression during the cell cycle.

## Introduction

Transcription is a central regulatory step of gene expression. It is a highly dynamic, multi-step process critical to basal cellular function, specification of cell identity, and response to environmental and developmental stimuli. Transcription is comprised of multiple steps including recruitment of the transcription machinery, initiation, promoter-proximal pausing, elongation and termination. During mitosis, the genome undergoes large-scale changes which disrupt the cell’s transcriptional program. These changes must be reversed following cell division, and the cell’s gene expression program re-established. Recent genomics studies have begun to change our understanding of the transcription regulatory events surrounding mitosis (for review, see Palozola, Lerner, and Zaret, 2019; Pelham-Webb, Murphy, and Apostolou, 2020; Raccaud, and Suter, 2018). However, it is not clear which steps of the transcription cycle are differentially regulated as the cell progresses from mitosis to G1 and how this regulation contributes to the gene expression program of daughter cells.

The dramatic morphological changes observed during cell division were originally demonstrated to co-occur with a global repression of transcription (Prescott, and Bender, 1962; Taylor, 1960). The underlying mechanism of repression was later demonstrated to involve premature termination of nascent transcription (Rothe, Pehl, et al, 1992; Shermoen, and O’Farrell, 1991). Nuclear import of transcription termination factor TTF2 was shown to occur specifically during the onset of mitosis, providing a molecular mechanism for global transcriptional repression (Jiang, Liu, et al, 2004). Further studies have also implicated mitotic hyperphosphorylation of general transcription factors (GTFs) and RNA polymerase II (RNAPII) itself as general mechanisms of repression (Bellier, Chastant, et al, 1997; Leresche, Wolf, and Gottesfeld, 1996; Segil, Guermah, et al, 1996; White, Gottlieb, et al, 1995; Zawel, Lu, et al, 1993). Recent genome-wide approaches, however, have begun to document low levels of active RNA polymerase during mitosis (Kang, Shokhirev, et al, 2020; Liang, Woodfin, et al, 2015; Palozola, Liu, et al, 2017). While the overall activity level of RNAPII during mitosis appears to be low, its role during mitosis remains unclear.

Despite hypercompaction of the genome during mitosis, many regulatory sites remain accessible (Gazit, Cedar, et al, 1982; Groudine, and Weintraub, 1982; Kadauke, Udugama, et al, 2012; Kuo, 1982; Martinez-Balbas, Dey, et al, 1995; Michelotti, Sanford, and Levens, 1997). Genome-wide approaches have recently demonstrated that retention of DNaseI-sensitivity in mitotic cells is a widespread phenomenon, but occurs preferentially at promoters rather than distal regulatory elements (Hsiung, Morrissey, et al, 2015). These observations have fueled the hunt for the underlying mechanisms, termed mitotic bookmarking, that maintain accessibility during mitosis (Michelotti, Sanford, and Levens, 1997). Initially, mitotic phosphorylation of sequence-specific transcription factors (TFs) was demonstrated to inhibit their association with DNA (Martinez-Balbas, Dey, et al, 1995; Segil, Roberts, and Heintz, 1991; White, Gottlieb, et al, 1995). Most TFs were shown to be excluded from chromatin during mitosis along with RNAP II (Hershkovitz, and Riggs, 1997; Kim, Du, et al, 1997; Parsons, and Spencer, 1997; Warren, Landolfi, et al, 1992). More recently, however, mitotic retention of TFs and histone modifications have been implicated in maintenance of the cell’s transcriptional program. One of the most well-studied bookmarking mechanisms involves the TBP/TFIID complex. TBP was recently shown to be maintained at a subset of interphase-loci during mitosis, along with transcriptionally-engaged RNAPII (Teves, An, et al, 2018). In contrast, numerous TFs were also shown to bind mitotic chromatin, but in a highly-dynamic, transcription-independent manner (Teves, An, et al, 2016). Acetylation of histones also appears to be an important mark for the rapid reactivation of specific genes following mitosis (Behera, Stonestrom, et al, 2019; Liu, Chen, et al, 2017; Liu, Pelham-Webb, et al, 2017; Pelham-Webb, Polyzos, et al, 2021). Recent results have shown that while BRD4 (a factor known to recognize histone acetylation) may be present on acetylated histones during mitosis, it is dispensable for gene reactivation (Behera, Stonestrom, et al, 2019). Overall, these studies suggest that while there is a global repression of transcription during mitosis, it appears that many transcription regulatory mechanisms continue to operate in a highly dynamic state.

The application of nascent RNA sequencing (for review of these methods, see Wissink, Vihervaara, et al, 2019) has also provided new insight into genome reactivation following mitosis. Waves of gene reactivation were demonstrated, beginning with genes involved in housekeeping functions and rebuilding the cell, while cell type-specific genes reactivated later (Kang, Shokhirev, et al, 2020; Palozola, Donahue, et al, 2017). Profiling of RNAPII with ChIP-seq in mice demonstrated a burst of transcription immediately following mitosis (Hsiung, Bartman, et al, 2016). Interestingly, functional tests of proposed mitotic bookmarks during global transcription reactivation have demonstrated that TBP and histone acetylation are only partially responsible for the re-establishment of transcription (Pelham-Webb, Polyzos, et al, 2021; Teves, An, et al, 2018). This suggests that multiple factors likely play a role in genome reactivation; some maintained during mitosis, while others must first be re-established before transcription resumes. Here we use Precision Run-On and sequencing (PRO-seq) to capture distinct transcription profiles during the cell cycle and uncover the contributing regulatory mechanisms. Integration of multiple genomic data sets during mitosis or upon mitotic exit show that all phases of transcription are dynamically regulated during this cellular transition with potentially independent consequences on transcription memory, the timing and amplitude of transcription reactivation, and the chromatin environment.

## RESULTS

### A distinct landscape of gene activity during mitosis

To document transcription and its regulation during mitosis, we employed PRO-seq on Hela-S3 cells following a thymidine-nocodazole block (Mahat, Kwak, et al, 2016; Whitfield, Sherlock, et al, 2002). This produced a highly enriched (>97%) population of cells arrested in prometaphase based on both FACS analysis and microscopy (Figure 1A, B). Because prometaphase cells have disassembled their nuclear envelope, we gently permeabilized the cell membrane rather than isolating nuclei prior to performing PRO-seq (Pelham-Webb, Polyzos, et al, 2021; Methods). Also, to account for global changes in transcription, fixed amounts of *Drosophila* S2 nuclei were added during the nuclear run-on step. When compared to asynchronous cells, we find a >30-fold reduction of transcription within gene bodies during mitosis (Figure 1C). While the majority of genes appear to be silenced, there is evidence of mitotic transcriptional activity at select genes (Figure 1D). In order to identify mitotic-active genes with confidence, we filtered the data for genes that maintain both read density and read coverage in gene bodies relative to asynchronous cells, resulting in 161 mitotic-active genes (Figures 1E and S1A, Methods). Activity of these genes during mitosis is also supported by Pol II ChIP-seq, nascent-seq, and an enrichment for active histone modifications in mitotic HeLa cells (Liang, Woodfin, et al, 2015, Figure S1B, C). Interestingly, mitotic-active genes have an increase in both promoter and gene body signal, but a decreased pausing index compared to inactive genes (Figure 1F and S1D). These results indicate that regulation of transcription at these genes during mitosis occurs, in part, through increased escape from promoter-proximal pausing and further demonstrates that the mitotic transcription signal is not a byproduct of residual non-synchronized cells.

**Figure 1.**
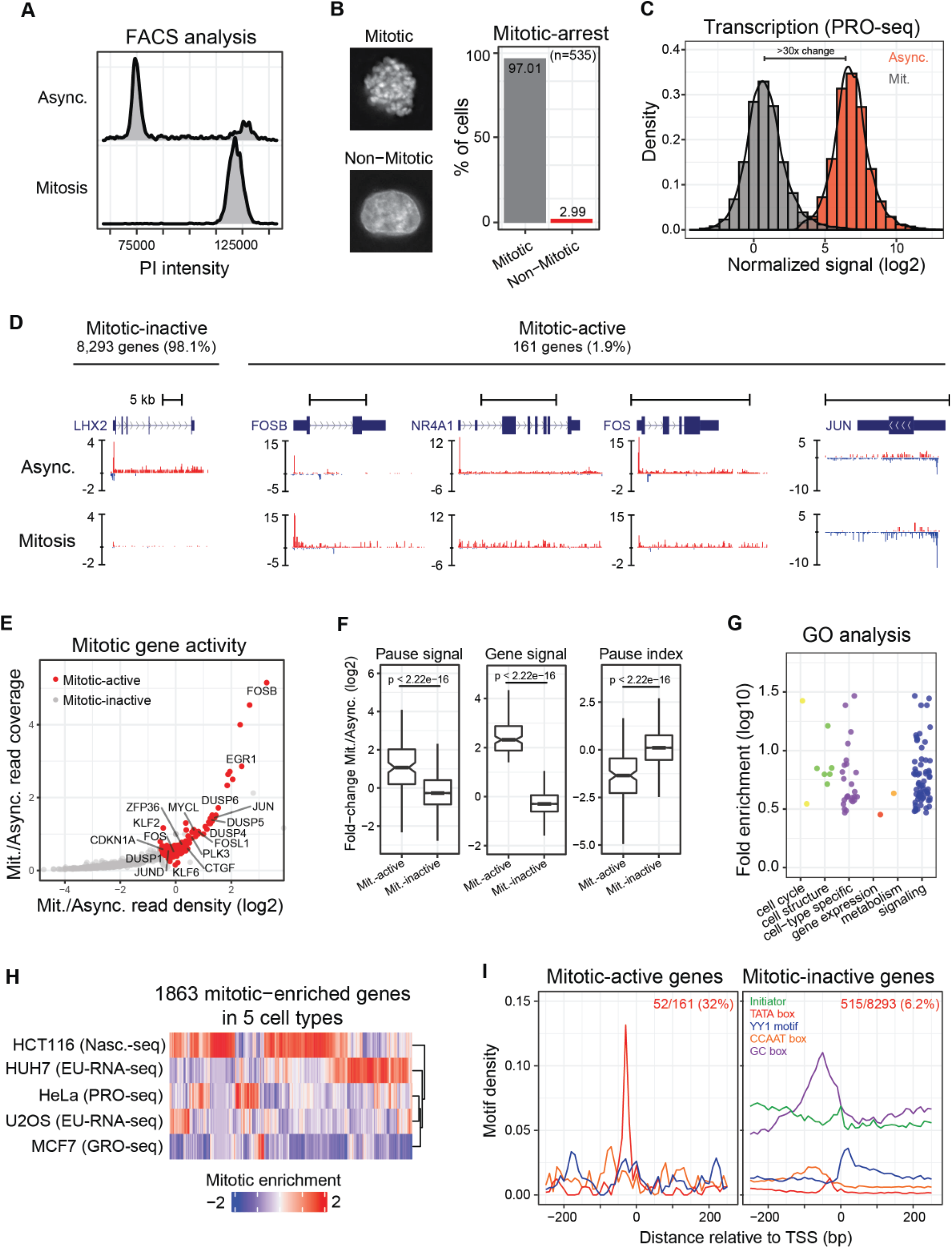
Gene activity and cell type-specific transcription profiles during mitosis. **(A)** FACS analysis of DNA content in asynchronous and mitotic HeLa-S3 cells stained with propidium iodide. Mitotic cells were arrested using thymidine-nocodazole block. **(B)** Representative images of cells following thymidine-nocodazole block following staining with DAPI (left panel). Quantification of mitotic and non-mitotic cells after thymidine-nocodazole block based on DNA morphology (right panel). (**C)** Density plot comparing spike-in normalized global transcription levels of protein coding and lincRNA genes in asynchronous and mitotic cells. **(D)** Genome browser tracks of PRO-seq signal at representative genes repressed or active during mitosis. All scale bars are 5kb and the Y-axis is spike-in normalized reads per million (RPM). **(E)** Scatterplot of the Mitotic/Async. ratio of read coverage and read density (log2) in gene bodies. Genes that maintain both normalized read density and coverage are identified as active (red) during mitosis. Select immediate-early genes are labeled. **(F)** Boxplots of the Mitotic/Async. fold-change in normalized PRO-seq signal at pause sites and gene bodies (left two panels). Boxplot of the Mitotic/Async. fold-change in pause index (right panel). P-values were calculated using the two-sided Wilcoxson’s rank sum test in R. **(G)** Gene ontology enrichment analysis of mitotic-active genes relative to terms associated with active genes in asynchronous cells. Individual GO terms were manually grouped into indicated classes (methods). **(H)** Heatmap of Mitotic/Async. differential expression analysis in five different cell types. **(I)** Composite profile of promoter sequence motifs in promoter regions of mitotic-active and mitotic-inactive genes.

When compared to all other genes, those active in mitosis are shorter than average and have a low expression level in asynchronous cells (Figure S1E). Gene ontology (GO) analysis of these mitotic-active genes demonstrated an enrichment of cell-signaling and cell-type specific processes (Figure 1G, Table S1). This included numerous transcription factors from the AP-1 family (FOS, JUN, ATF, and MAF) as well as other members of the “early-response genes” (Figure 1E). This group of genes has been implicated in a number of cellular processes from cell growth and proliferation, to stress responses, are often dysregulated in cancers, and are thought to be co-regulated (Vacca, Itoh, et al, 2018). We then compared mitotic transcriptional activity across several cell types during mitosis by incorporating nascent transcription data from previous studies (Kang, Shokhirev, et al, 2020; Liang, Woodfin, et al, 2015; Liu, Chen, et al, 2017; Palozola, Donahue, et al, 2017). While we detect mitotic enrichment for some genes across all five cell types (eg. FOSB and NR4A3), many genes are enriched during mitosis in a cell type-dependent manner (Figure 1H, S1B, F-I). This includes genes associated with the GO-term “cell differentiation” and other cell-type specific terms, although the annotated functions for the cell-specific genes do not necessarily correspond to each individual cell type. However, GO analysis of mitotic-enriched genes across all five cell types revealed a weak enrichment for terms associated with epithelial cells (Figure S1J). Interestingly, these genes also demonstrated enrichment for various pathways related to cell cycle control and cancer (Figure S1K).

Further evidence for distinct regulation of transcription during mitosis comes from analysis of core promoter motifs. While most human promoters lack a TATA element, we find overrepresentation (>5-fold enrichment) of this regulatory sequence in promoters of our mitotic-active genes (Figure 1I, S1L). This suggests that direct binding of TBP to promoters may be critical for the maintenance of transcription during mitosis and is supported by live cell imaging of both TBP and Pol II in mitotic mouse ESCs (Teves, An, et al, 2018). Overall, these findings suggest that the genes remaining active during mitosis exhibit cell type-specificity and require strong core promoter elements.

### Promoter activity and maintenance of the local chromatin environment during mitosis

Previous studies have demonstrated that while chromatin is highly compacted during mitosis, promoters preferentially remain accessible (Hsiung, Morrissey, et al, 2015; Kadauke, Udugama, et al, 2012; Teves, An, et al, 2016). To compare chromatin accessibility with our PRO-seq data, we incorporated deeply sequenced mitotic ATAC-seq from another study (Oomen, Hansen, et al, 2019). We observe a global retention of accessibility as mitotic active and inactive genes have relatively little difference in the level of chromatin accessibility during mitosis (Figure S1C, S2A). However, despite the overall reduction in transcription during mitosis, PRO-seq signal is preferentially retained at sites of transcription initiation compared to gene body regions (Figure S2B). Given this partial mitotic retention of Pol II at promoters, we further explored the interplay between local chromatin architecture and transcription during mitosis. We created a comprehensive map of genomic regulatory elements by incorporating information from ATAC-seq, PRO-seq, and ChIP-seq data (Figure S2C, Methods). We find that most promoters (98%) and 18% of enhancers appear to remain transcriptionally active during mitosis (Methods). Notably, both of these groups also maintain significant levels of accessibility during mitosis (Figure 2A). This is in stark contrast to the loss of accessibility at transcriptionally inactive enhancers such as poised enhancers, CTCF binding sites (structural regulatory elements), and enhancers that are transcriptionally repressed during mitosis (Figure 2A). Grouping all active enhancers by presence/absence of mitotic PRO-seq signal demonstrated that chromatin accessibility during mitosis is tightly associated with transcription activity, regardless of enhancer strength in asynchronous cells (Figure 2B). These results suggest that during mitosis, continued transcription initiation is associated with maintenance of chromatin accessibility,

**Figure 2.**
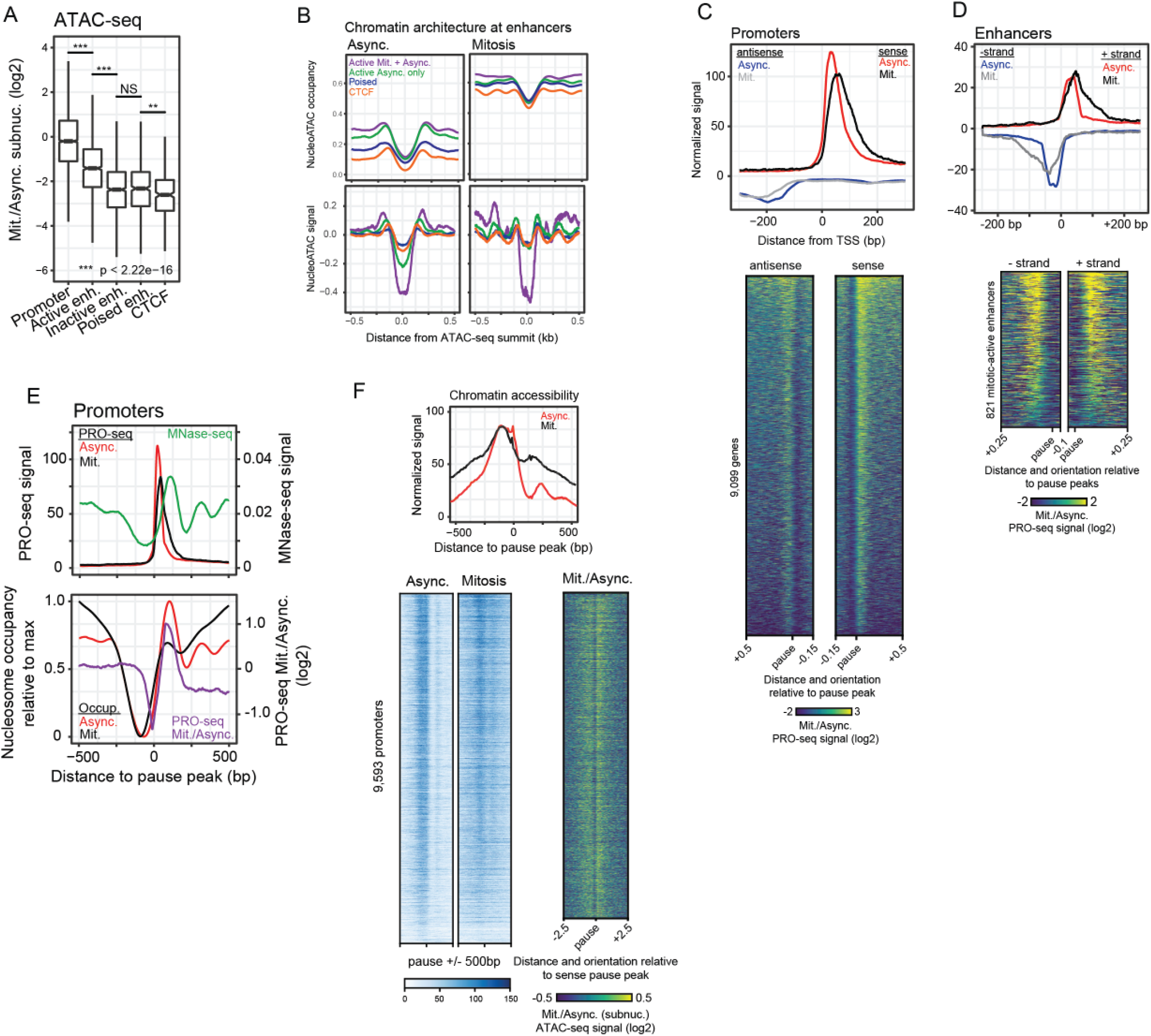
Transcription and chromatin accessibility during mitosis. **(A)** Boxplots of normalized ATAC-seq signal at genomic regulatory elements. Enhancers are classified by their transcriptional activity during mitosis. **(B)** Nucleosome occupancy (top panels) and nucleosome signal (bottom panels) profiles at genomic regulatory elements (excluding promoters) in mitotic and asynchronous cells. **(C)** Average profiles of normalized PRO-seq signal relative to TSSs in mitotic and asynchronous cells (upper panel). Heatmaps of log2 fold-change Mitosis/Async. PRO-seq signal (lower panels) relative to both the antisense pause peak (left heatmap) and the sense pause peak (right heatmap). **(D)** Average profiles of normalized PRO-seq signal relative to enhancer centers in mitotic and asynchronous cells (upper panel). Heatmaps of log2 fold-change Mitosis/Async. PRO-seq signal (lower panels) relative to minus-strand pause sites (left heatmap) and plus-strand pause sites (right heatmap) of enhancers. (**E)** Average profiles of normalized PRO-seq and MNase-seq signal relative to the pause sites (top panel). Average profile of estimated nucleosome occupancy by nucleoATAC in async. and mitotic cells and log2 fold-change Mitotic/Async. PRO-seq signal relative to the pause sites (lower panel). **(F)** Average profile of normalized ATAC-seq signal relative to sense pause sites in asynchronous and mitotic cells (top panel). Heatmaps of normalized ATAC-seq signal in asynchronous and mitotic cells (bottom left two panels) and log2 fold-change of Mitotic/Async. ATAC-seq signal (bottom right panel).

We then closely examined the PRO-seq signal at promoters for changes in behavior during mitosis. Composite profiles centered on promoter-proximal pause sites revealed that the mitotic-retained polymerase shifts to a position ~21bp (+56bp) downstream of the normal pausing site at promoters and enhancers. (Figure 2C-E S2D). Interestingly, this shift in pause signal closely resembles that seen in NELF-depleted cells (Aoi, Smith, et al, 2020). To investigate whether the altered pausing pattern plays a role in maintenance of Chromatin accessibility during mitosis, we compared pausing profiles to surrounding nucleosome occupancy. MNase-seq data from asynchronous cells shows a peak position of +124 bp for the +1 nucleosome, putting it close enough to be in contact with the downstream shifted polymerase (Figure 2E, upper panel, Kfir et al., 2015). We then used NucleoATAC to compare changes in local chromatin architecture of mitotic and asynchronous cells. (Oomen, Hansen, et al, 2019; Schep, Buenrostro, et al, 2015). In asynchronous cells, as expected, the nucleosome occupancy profile from NucleoATAC is highly similar to the MNase-seq profile with the +1 nucleosome showing the highest occupancy compared to the surrounding chromatin (Figure 2E, compare top and bottom panels). During mitosis, promoters maintain low nucleosome occupancy and the +1 nucleosome can be detected, however, +1 nucleosome occupancy is decreased compared to the surrounding chromatin (Figure 2E, bottom panel). This suggests that although there is a global increase in nucleosome occupancy across genes during mitosis, there is a mechanism in place that maintains an open promoter while competing with nucleosome deposition. Intriguingly, the greatest change in PRO-seq density during mitosis directly overlaps the +1 nucleosome, suggesting a potential clash between Pol II with the +1 nucleosome during mitosis (Figure 2E, bottom panel). In support of this, we also detect an altered ATAC-seq profile and an increase in subnucleosomal reads in the ATAC-seq data that overlap promoter-flanking nucleosomes indicating that these nucleosomes are being moved or destabilized during mitosis. (Figure 2F). Altogether the data indicate that while most transcriptional activity is repressed during mitosis, both promoters and select enhancers continue to undergo initiation as well as a distinct mode of pausing, and that this activity is associated with maintenance of chromatin accessibility during mitosis.

### Genome reactivation following mitosis is highly dynamic

To measure the kinetics of transcription reactivation following mitosis we performed PRO-seq on cells in a time course up to 12hr after release from nocodazole. Cellular DNA content was monitored with FACS, demonstrating progression through cytokinesis at 1.5-2hr post-release and entry into S-phase beginning at 12hr (Figure 3A). Gene reactivation begins between 1 and 1.5hr following mitosis which is consistent with results from Pol II ChIP-seq (Hsiung, Bartman, et al, 2016) and pulse labeling of RNA (Palozola, Donahue, et al, 2017) (Figure 3B). Additionally, high-resolution nascent RNA profiling methods such as PRO-seq allow the initial waves of polymerase to be detected as it progresses into gene bodies (Figure 3C, D). This feature can be used to derive a transcription rate of RNA polymerase by capturing the leading edge of transcription in successive time points (Figure 3D, Danko, Hyland, et al, 2015). Doing so, we find polymerase traveling rates of 2 - 2.5kb/min as well as a slight increase in rate as the wave progresses further into the gene body (Figure 3E). This is in agreement with previous findings (Danko, Hyland, et al, 2015; Jonkers, Kwak, and Lis, 2014) and suggests that bursts of transcription reactivation detected upon exit from mitosis (Hsiung, Morrissey, et al, 2015) are not due to increased transcription rates. To validate the timing of transcriptional reactivation and cellular functions, we examined gene families known to be tightly regulated during the cell cycle. As expected, the core histone genes were specifically reactivated at 8-12hr post-release, coinciding with the onset of S-phase (Figure 3F, left panel). Cyclin genes have long been known to control cellular progression through interphase. We found their timing of expression matches their known functions in the various cell cycle stages (Figure 3F, right panel). Also, further analysis revealed that cyclin genes primarily involved in transcription werereactivated earlier than cyclins with a primary function in cell cycle control (Figure 3F). Whereas our data appears to show faster kinetics than previous studies that examine mRNA (Whitfield, Sherlock, et al, 2002) (Figure S3A), the order of gene activation is consistent. The difference in apparent timing between studies is expected since PRO-seq provides an immediate and direct measure of transcription changes and is much more sensitive than microarrays.

**Figure 3.**
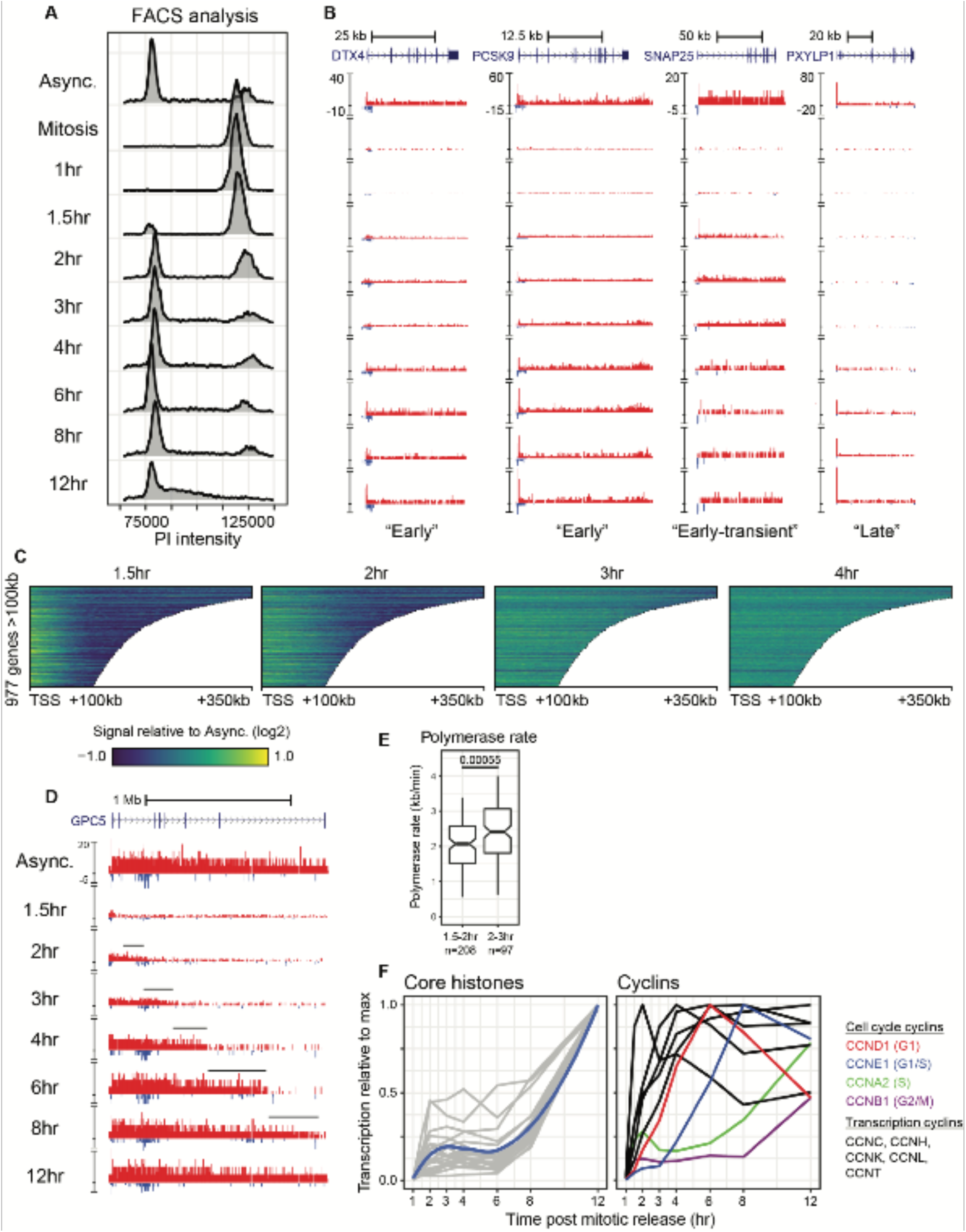
PRO-seq provides a high-resolution measurement of genome reactivation following mitosis. **(A)** FACS analysis of DNA content in HeLa-S3 cells stained with propidium iodide following mitotic arrest and release. **(B)** Browser tracks of genes representative of different reactivation kinetics following mitosis. Y-axis is spike-in normalized signal. **(C)** Heatmaps of log2 fold-change PRO-seq signal during transcription reactivation at genes >100kb in length. **(D)** Browser tracks of a representative long gene demonstrating the initial wave of transcription reactivation. Y-axis is spike-in normalized signal. **(E)** Boxplots of the calculated RNA polymerase rates in genes. **(F)** Reactivation kinetics of gene families known to be associated with the cell cycle. Y-axis is spike-in normalized signal relative to maximum transcription level during the time course.

### Gene reactivation classes are associated with distinct cellular functions

To assess kinetics of reactivation genome-wide, we grouped genes by the timepoint at which they reach half-maximum signal (Figures 4A, B and S4A, Methods). In order to detect rapid changes in transcription and eliminate the effect of varying gene lengths on measurements of reactivation timing, genes were trimmed to a maximum length of 10kb. We find 28% of genes are reactivated within the first 2hr and subsequently 82% of genes are reactivated by four hours post mitotic-release (Figure S4B). Importantly, we do not detect biases in our kinetic classes toward short gene length or high expression level in asynchronous cells in the early activated genes (Figure S4C, D). Gene ontology (GO) analysis of each kinetic class, revealed an enrichment in distinct cellular functions. The earliest class of genes demonstrate overrepresentation of cell structure-related functions as well as cellular signaling. (Figure 4C). Subsequent waves of reactivation are enriched for genes involved in both cellular metabolism and gene expression at 3-4hr, followed by cell type-specific functions at 6-8hr (Figure 4C). As expected, the 12hr class is highly enriched for genes involved in the cell cycle, consistent with the onset of S-phase (Figure 4C). While the majority of genes reactivate in early-G1, we detect distinct functional waves of activation following mitosis as described in previous studies (Palozola, Donahue, et al, 2017).

**Figure 4.**
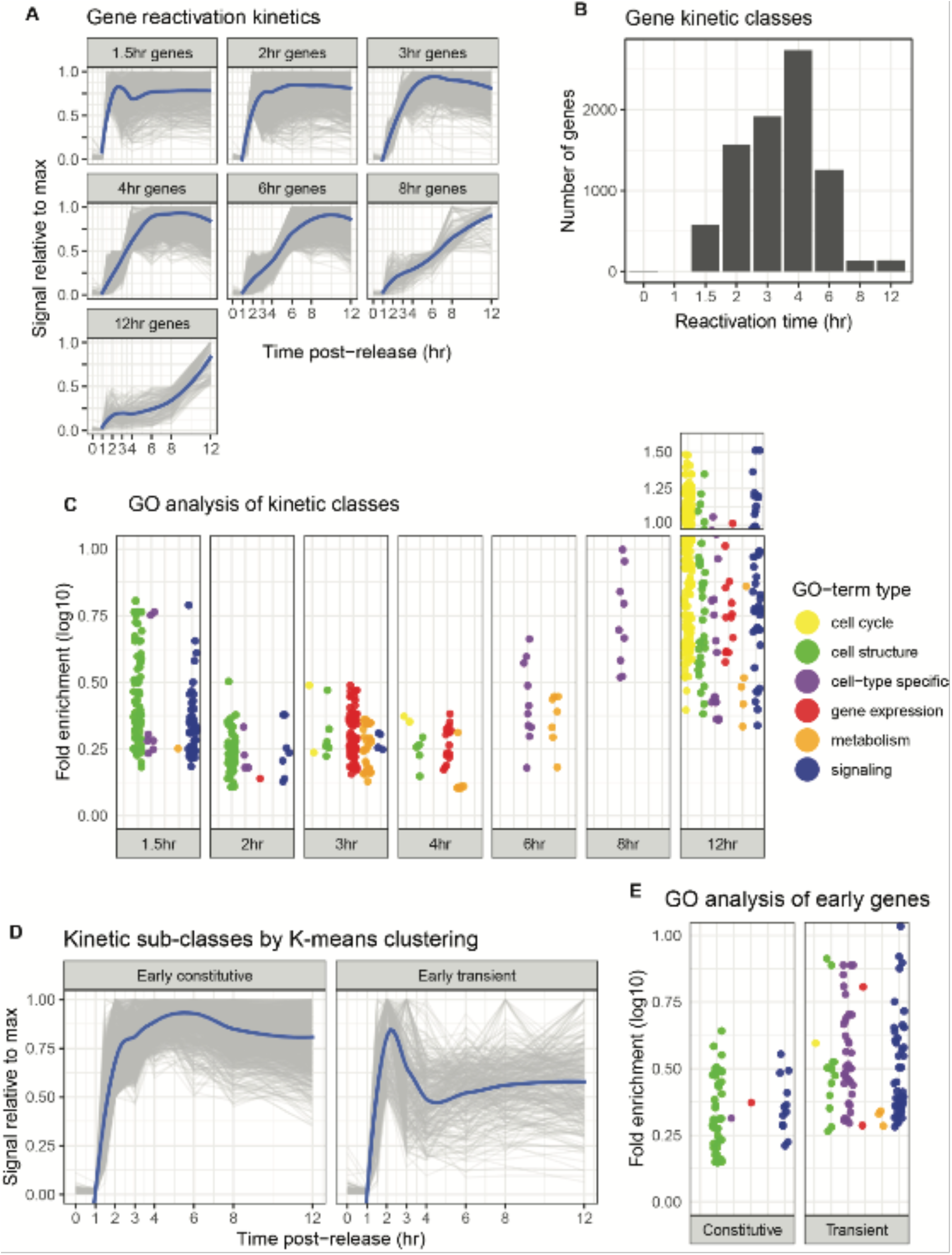
Functional gene groups have different timing and duration of reactivation during cell cycle progression. **(A)** Line plots of gene kinetics during the 12hrs after release from mitosis for each kinetic class. A summary line (blue) was generated for each kinetic class by fitting a loess curve of the 1-12hr timepoints. Y-axis is spike-in normalized signal relative to maximum transcription level throughout the time course. **(B)** Summary bar plots showing the number of genes in each kinetic class. **(C)** Gene ontology enrichment analysis of each gene kinetic class relative to terms associated with all active genes throughout the time course. Individual GO terms were manually grouped into indicated classes (methods). **(D)** Line plots of gene kinetics during the 12hrs after release from mitosis for early genes clustered by signal after initial reactivation. Y-axis is spike-in normalized signal relative to maximum transcription level throughout the time course. **(E)** Gene ontology enrichment analysis of early genes clusters relative to terms associated with all active genes throughout the time course.

The order in which genes are reactivated after mitosis has been a focal point in studies of cell-type specific gene expression. Central to this question is whether early activation of genes that specify cell type is critical to maintenance of cell identity (Kadauke, Udugama, et al, 2012). Our initial results are similar to those found in both HUH7 hepatoma cells and U2OS osteosarcoma cells (Kang, Shokhirev, et al, 2020; Palozola, Donahue, et al, 2017), that suggest genes responsible for cell growth are activated first, followed by cell-type specific genes (enrichment of GO-terms related to “development”, “differentiation”, “morphogenesis”). However, these earliest activated gene classes (1.5hr and 2hr) appear to have a high degree of variability in their expression levels following initial activation (Figure 4A). We therefore combined these groups and performed k-means clustering, leading to distinct classes of early genes based on overall kinetics throughout the time course, which we term “early-constitutive” and “early-transient” (Figure 4D). GO analysis of these classes demonstrate that early-constitutive genes are associated with cell structure and housekeeping functions, while the early transient genes are enriched in signaling and cell type-specific roles (Figure 4E). In addition to timing of activation, robust transcription of genes is hypothesized to be a contributor to cell-specific gene expression. A recent study in mouse erythroblasts suggested that gene reactivation following mitosis often occurs in a robust burst of transcription (Hsiung, Bartman, et al, 2016). Using our data, we identified 365 “burst” genes whose spike-in normalized expression levels during the time course are greater than expression levels in asynchronous cells (Figure S4E, Methods). While this gene class was enriched in both cell structure and signaling-related functions, we found early-genes that did not burst were enriched in functions related to metabolism and gene expression (Figure S4F). Overall, our data suggests that functional gene classes are not only controlled with respect to timing of reactivation, but also amplitude and persistence through interphase. We find a group of rapidly activated, but transiently expressed genes that appear to function in cell-type specific processes, potentially playing a role in the maintenance of cell identity.

### Enhancer reactivation following mitosis is linked to functional waves of gene reactivation

PRO-seq data allows for the identification of enhancers based on a bidirectional transcription signature similar to that of promoters (Core, Martins, et al, 2014; Danko, Hyland, et al, 2015; Methods). The production of nascent RNA at enhancers is a proxy for enhancer activity (Henriques, Scruggs, et al, 2018; Mikhaylichenko, Bondarenko, et al, 2018; Tippens, Liang, et al, 2020) and therefore allows us to directly compare gene and enhancer kinetics following mitosis. Overall, we find enhancer reactivation appears to be delayed relative to gene bodies (Figure 5A). Grouping enhancers into kinetic classes by signal relative to max shows that the majority of enhancers (68%) are reactivated in mid- to late-G1(6-8hrs post-mitosis) (Figure S5A, B). Interestingly, grouping enhancers by distance to the nearest promoter revealed a relationship between the timing of enhancer reactivation and gene-proximity (Figure 5B). GO analysis of enhancer classes using distance-based gene assignments revealed waves of functional enrichment that matched those of gene reactivation groups (McLean, Bristor, et al, 2010) (Figure 5C). As a more direct test of this association, we assigned enhancers to the nearest gene and compared their kinetic classes (Figure S5C, Methods). The profiles of enrichment suggest that many gene-enhancer pairs are reactivated together, with enrichment of matching kinetic classes found more often than expected by random assignments (Figure 5D). To further validate these findings, we performed ATAC-seq on our asynchronous, mitotic, 2hr and 6hr timepoints. Analysis of this data confirmed a delay in enhancer reactivation relative to promoters and also revealed a delay in accessibility of other regulatory elements including poised enhancers and CTCF sites (Figures 5E, 5F). While our analyses suggest an overall delayed reactivation of enhancers, these measurements may be due, in part, to altered pause-escape rates during early-G1 (see below). Nonetheless, we are able to detect a link between early-activated gene-enhancer pairs.

**Figure 5.**
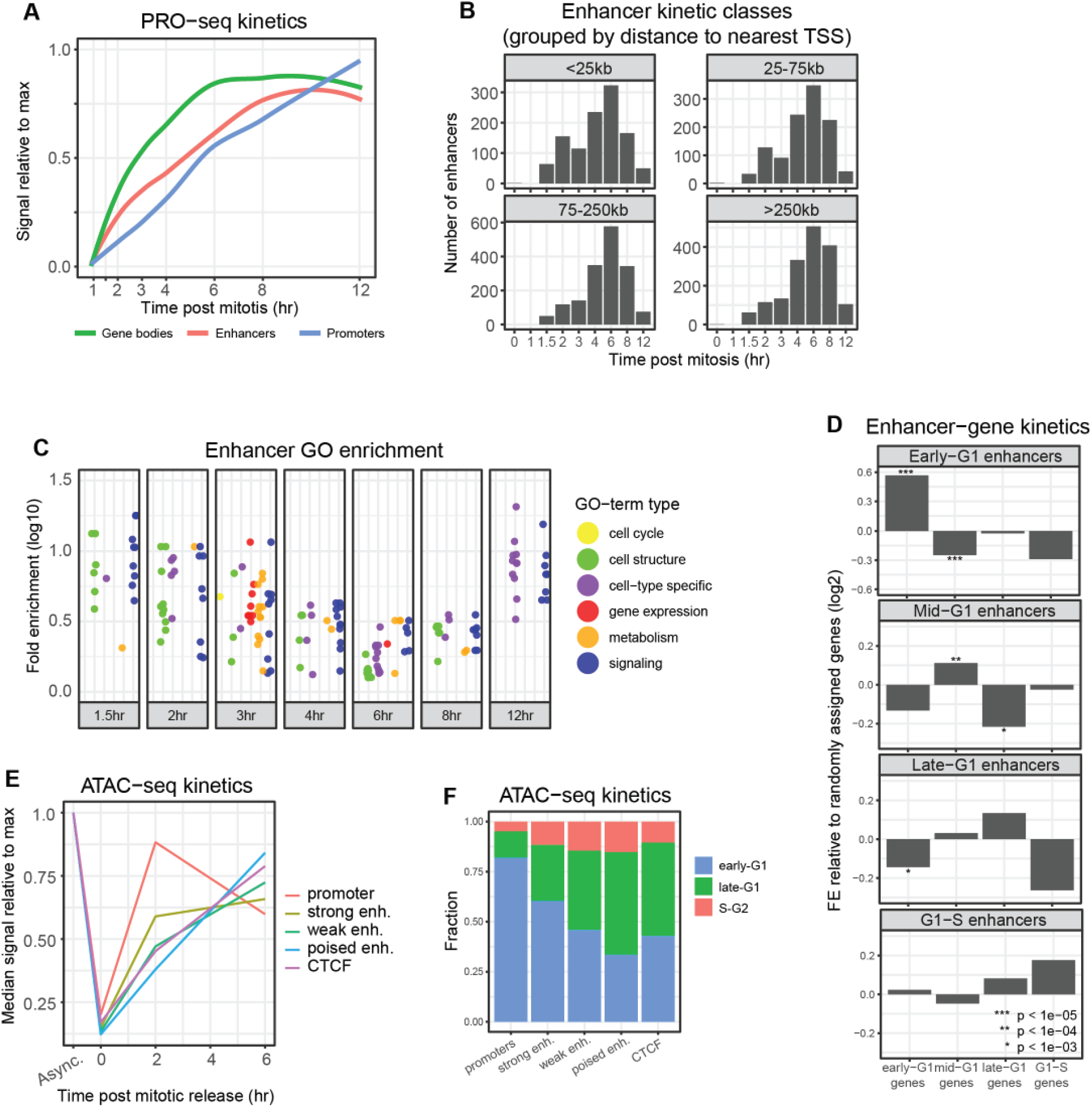
Kinetics of post-mitotic enhancer reactivation. **(A)** Summary line plots of reactivation kinetics of all genes compared to all transcriptionally active enhancers. Y-axis is spike-in normalized signal relative to maximum transcription level throughout the time course. **(B)** Summary bar plots of the number of enhancers in each kinetic class grouped by distance to the nearest active gene TSS. **(C)** Gene ontology enrichment analysis of enhancers when assigned to nearby genes. Individual GO terms were manually grouped into indicated classes (methods). **(D)** Enrichment of gene kinetic classes when assigned to each enhancer by reactivation time. Enhancers are assigned to the nearest TSS of each gene kinetic group and enrichment is calculated by comparing to random gene assignments. **(E)** Summary line plot of mean reactivation kinetics of regulatory elements measured by ATAC-seq. Y-axis is normalized signal relative to maximum accessibility level throughout the ATAC-seq time course. **(F)** Summary bar plot of reactivation timing for regulatory element classes.

### A role for mitotic transcriptional activity in promoter bookmarking and gene reactivation kinetics

The propagation of transcriptional states through cell division was initially characterized by the maintenance of DNase hypersensitive sites at individual loci (Gazit, Cedar, et al, 1982; Groudine, and Weintraub, 1982; Kadauke, Udugama, et al, 2012; Martinez-Balbas, Dey, et al, 1995; Michelotti, Sanford, and Levens, 1997). Genome-wide approaches have revealed this to be a common feature among regulatory elements, occurring with variable degrees of accessibility during mitosis (Hsiung, Morrissey, et al, 2015; Oomen, Hansen, et al, 2019; Owens, Papadopoulou, et al, 2019; Pelham-Webb, Polyzos, et al, 2021; Teves, An, et al, 2016). Numerous factors have been implicated in the maintenance of mitotic-chromatin accessibility, including a variety of histone modifications and protein factors (Behera, Stonestrom, et al, 2019; Javasky, Shamir, et al, 2018; Pelham-Webb, Polyzos, et al, 2021; Teves, An, et al, 2018). This phenomenon, termed “mitotic bookmarking,” has been hypothesized to both maintain the cell’s transcriptional program and control the kinetics of gene reactivation following mitosis (Hsiung et al., 2015). To investigate the effect of various potential bookmarks on gene reactivation, we integrated our PRO-seq data with recently published ChIP-seq and CUT&RUN data in mitotic HeLa cells (Javasky, Shamir, et al, 2018; Liang, Woodfin, et al, 2015; Oomen, Hansen, et al, 2019). Both mitotic H3K9ac and H3K4me3 marks are enriched in early- and mid-G1 gene promoters (accounting for >75% of genes) (Figure 6A). While there is an overall reduction in H3K27ac signal during mitosis, this mark is also preferentially retained at early- and mid-G1 gene promoters compared to late-G1 (Figure 6 A). Interestingly, measurements of promoter accessibility and RNA polymerase occupancy during mitosis exhibit the strongest relationship with early reactivation kinetics (Figure 6A) (Liang, Woodfin, et al, 2015; Oomen, Hansen, et al, 2019). In a reciprocal analysis to test whether these marks are predictive of gene reactivation, we compared gene kinetics of top and bottom quartiles for each potential bookmark (Figure 6B). Similar to the previous analysis, there is a noticeable difference in kinetics of these groups for all marks associated with transcription (H3K27ac, H3K9ac, H3K4me3) (Figure 6B, left panel). Measurements of Pol II activity during mitosis (both ChIP-seq and PRO-seq) were among the most predictive of rapid reactivation kinetics. Furthermore, Pol II ChIP-seq after inhibition of pause-release with flavopiridol in mitotic cells (Liang, Woodfin, et al, 2015) resulted in the strongest relationship with gene reactivation kinetics. Pairwise analyses of each potential bookmark also demonstrates a slight increase in predictive power of many factors (Figure 6B, right panel). Our results suggest that while individual bookmarks may ensure expression in the daughter cells, the timing of gene reactivation is linked more closely to the overall transcription activity and maintenance of promoter structure during mitosis.

**Figure 6.**
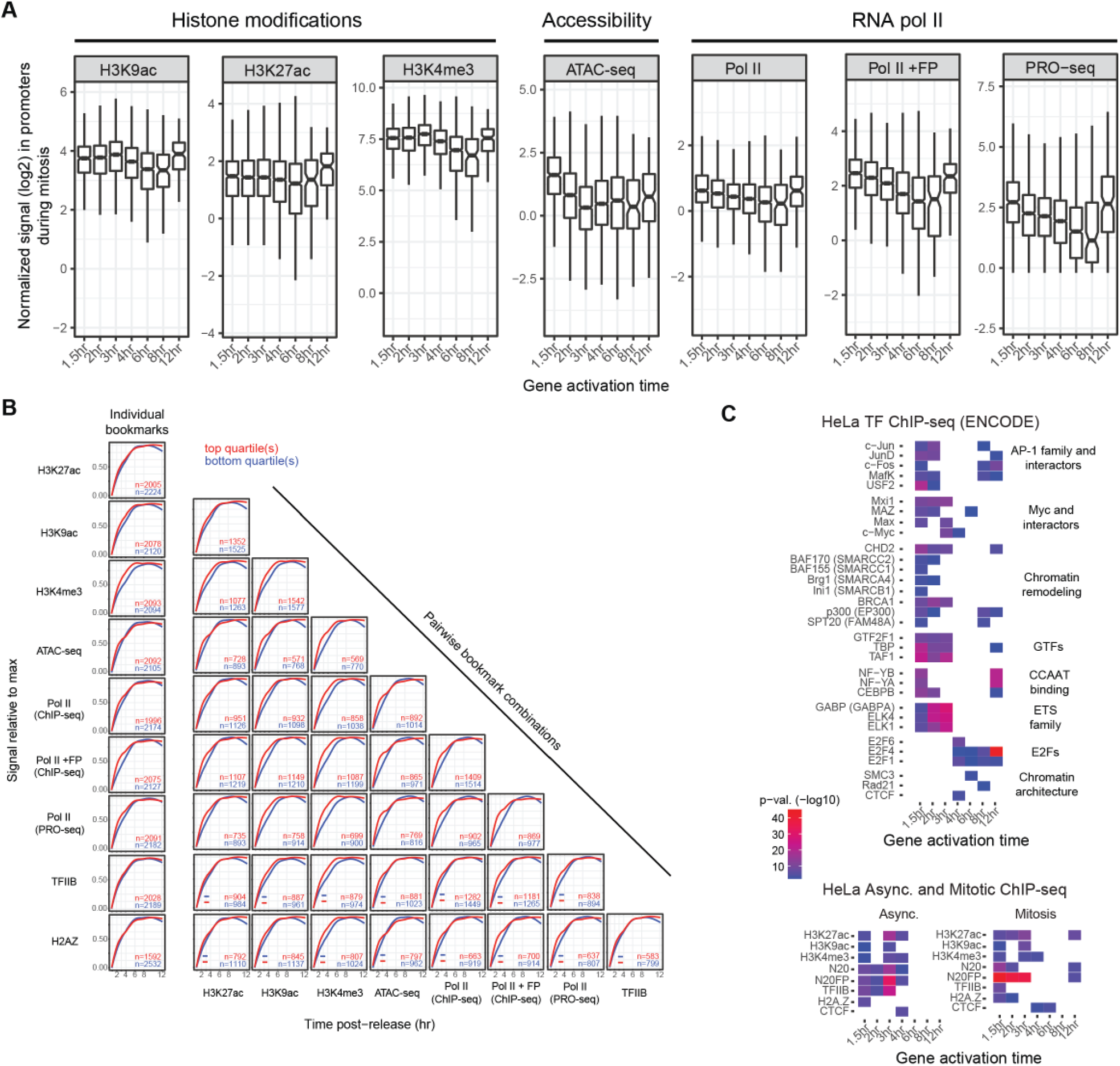
Diverse modes of molecular bookmarking contribute to gene reactivation following mitosis. **(A)** Boxplots of promoter signal during mitosis in each gene kinetic class. Y-axis is normalized reads per million (RPM). **(B)** Summary line plots of gene reactivation kinetics comparing top and bottom quartiles of potential bookmarking factors individually (far left column) and in pairs. Y-axis is spike-in normalized signal relative to maximum transcription level throughout the time course. **(C)** Enrichment of ENCODE transcription factors (top panel) and peaks from ChIP-seq data in asynchronous and mitotic cells (bottom panel) at promoters of each gene kinetic class.

Gene promoters are critical sites of signal integration, typically involving a host of GTFs as well as sequence-specific TFs that coordinate recruitment and initiation of Pol II. To begin to understand the role of such factors in gene reactivation following mitosis, we integrated the ENCODE TFBS database into our analyses. The earliest kinetic classes are most strongly enriched for members of the TFIID complex such as TBP and TAF1 as well as USF2 (a factor known to associate with members of the AP-1 family) while also showing weak enrichment for numerous factors involved in chromatin remodeling and Myc-interactors (Figure 6C). We detected enrichment of ETS family (ELK1 and ELK4) and GABPA transcription factors in 2-3hr kinetic classes. Factors in the E2F family are enriched in later kinetic classes, corresponding with their function in regulation of the cell cycle (Attwooll, Lazzerini Denchi, and Helin, 2004). Motif analysis supported the involvement of specific transcription factor families with distinct gene reactivation kinetics following mitosis. We detect a transition from the AP-1 and NF-Y motif families to the ETS and YY1 families during progression from early to mid G1 (Figure S6A, B). Motif analysis also revealed enrichment of LIN54 in the 12hr gene class, consistent with its function in the G2/M transition (Schmit et al., 2007). Overall, these results highlight the role of transcription factors in the control of gene reactivation following mitosis and suggest that certain factors cooperate with promoters to efficiently recruit the general transcription machinery at the earliest activated genes.

### Regulation through promoter-proximal pausing fine tunes transcription throughout the cell cycle

Intriguingly, many early-activating genes exhibit a lag in the re-establishment of PRO-seq signal at the promoter-proximal pause site relative to the gene body (Figure 3B, Figure 7A). Indeed, when compared to asynchronous cells, active genes during early- and mid-G1 have dramatically lower pausing indexes (Figure 7B). This suggests that following recruitment and initiation, RNA polymerase is more rapidly released from pausing during early-G1 when compared to asynchronous cells. Dynamic control of pausing was further supported by our analysis of early activated genes. Comparing the pausing index of these classes reveals that activation of early-transient genes is accompanied by low pausing levels, while their subsequent repression later in the cell cycle occurs concomitant with an increase in pausing levels (Figure S7A). This implicates pause-release as the central regulatory step in control of the early-transient gene class. Overall, these observations suggest that regulation through pausing is dynamic through the cell cycle and contributes to rapid reactivation of transcription after mitosis and fine-tuning of transcription over time.

**Figure 7.**
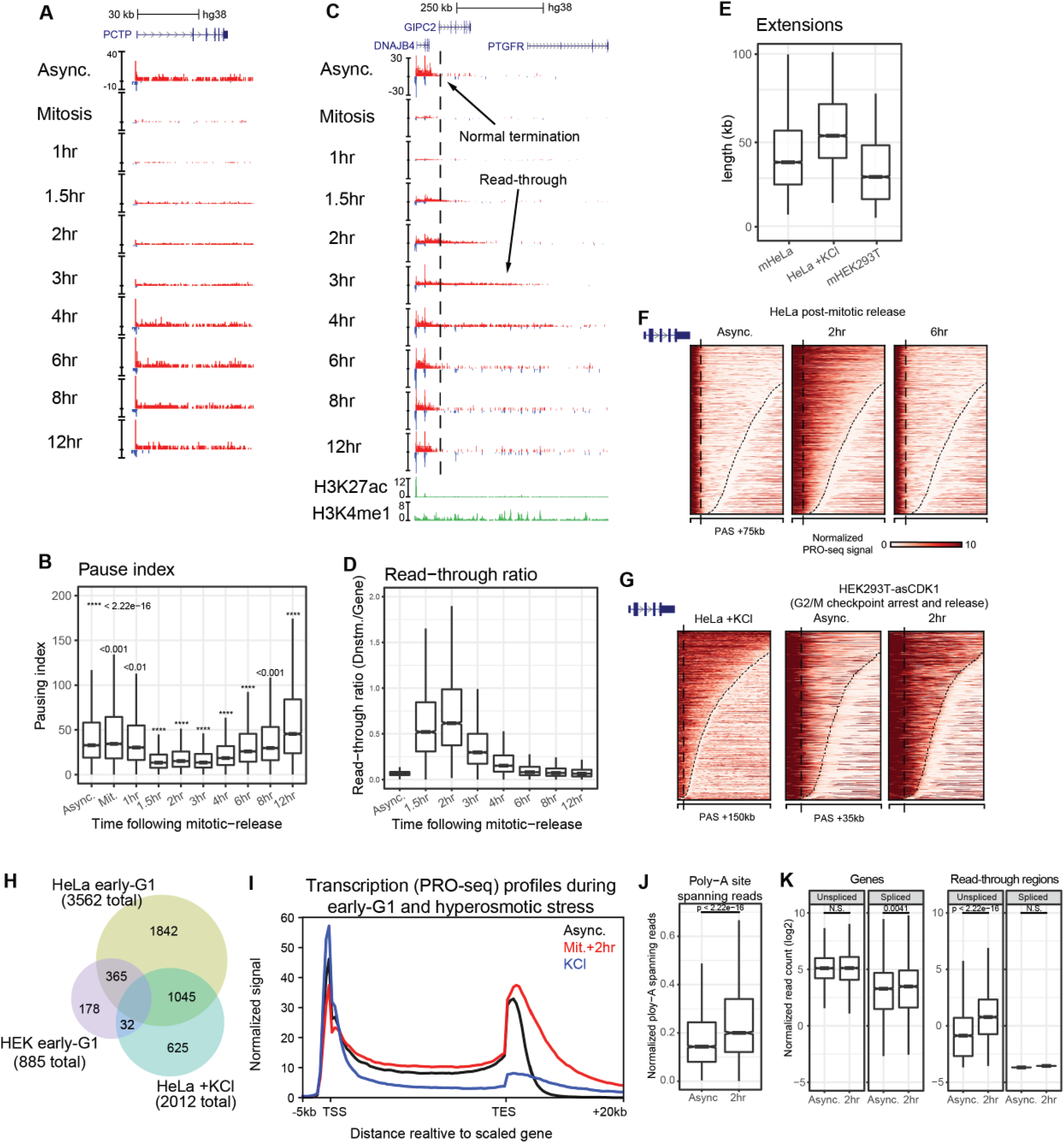
Transcription elongation phenotypes during genome reactivation. **(A)** Browser tracks of representative gene demonstrating a reduced pausing index immediately following mitosis. Y-axis is spike-in normalized PRO-seq signal. **(B)** Boxplot of pausing index of all active genes at each timepoint following release from mitosis. Each timepoint is compared to asynchronous cell using the two-sided Wilcoxson’s rank sum test function in R. **(C)** Representative browser image of a gene demonstrating read-through transcription immediately following release from mitosis including ChIP-seq data. Dashed line represents the end of the transcription termination zone in asynchronous cells. Y-axis is spike-in normalized PRO-seq signal and ChIP-seq reads per million (RPM). (**D)** Boxplots of read-through ratio (downstream read density divided by gene body read density) at all active genes during the time course. **(E)** Boxplot of read-through lengths in HeLa cells following mitotic release, HeLa cells undergoing hyperosmotic shock, and HEK293T-asCDK1 cells following release from the G2/M checkpoint. **(F)** Heatmap of downstream gene regions in HeLa cells undergoing read-through transcription following mitosis. Dashed lines represent the end of the transcription termination zone in asynchronous cells. **(G)** Heatmaps of downstream gene regions undergoing read-through transcription in HeLa cells following hyperosmotic shock (left panel) and HEK293T-asCDK1 cells following release from G2/M arrest. Dashed lines represent the end of the transcription termination zone in asynchronous cells. **(H)** Venn diagram comparing genes undergoing read-through transcription under various conditions and in different cell types. **(I)** Average profile of sense PRO-seq signal comparing gene signal in untreated asynchronous cells, cells following hyperosmotic shock, and 2hr after release from mitosis. **(J)** Boxplot of poly-A spanning reads from full-length PRO-seq data comparing asynchronous cells and those 2hr post-release from mitosis.

### Widespread transcription read-through creates a unique transcriptional landscape in early G1

One unexpected transcription phenotype we observed during gene reactivation is widespread read-through transcription specifically during early- and mid-G1 (Figure 7C, D). Recently, this phenomenon has been associated with various cell stressors and aberrant transcriptional complexes (Erickson, Sheridan, et al, 2018; Vilborg, Passarelli, et al, 2015). To test whether this was simply due to nocodazole-induced stress, we treated asynchronous cells for 2 hours at the concentration used in M-phase arrest (100ng/mL). While nocodazole had no apparent effect on transcription termination, hyperosmotic stress did induce read-through in a manner similar to recent work (Figure S7B) (Castelo-Branco, Amaral, et al, 2013; Vilborg, Passarelli, et al, 2015). Also, to corroborate the association of read-through transcription within the cell cycle, we generated an altered-specificity CDK1 (CDK1-as) cell line in HEK293T cells (Bishop, Ubersax, et al, 2000; Gravells, Tomita, et al, 2013). Treatment of these cells with 1-NM-PP1 for 20 hours induces an G2/M arrest by specifically inhibiting CDK1-as. Following washout of inhibitor, we collected cells at 2.5- and 7.5-hours post-release and performed PRO-seq. In these cells, we also detect read-through transcription specifically during early-G1 (Figure S7C). We also detect read-through transcription in nascent RNA from other cell types, including non-transformed cell lines, albeit with less intensity presumably due to the methods used, time-points chosen, and speed with which cells transition through this phase (Figure S7D, E). Assessment of read-through transcription during the mitotic time course revealed read-through transcription at 3,562 genes (64%), extending as far as 300kb (Figure 7E). In comparison, we detected read-through transcription at 2,012 genes (36%), and 885 (17%) following hyperosmotic shock or G2/M synchronization in HEK cells, respectively (Figure 7F-H). Notably, mitotic exit and osmotic stress conditions have opposite effects on transcriptional activity within genes (Figure 7I), and genes experiencing readthrough show moderate overlap between conditions or cell types. Altogether, these results show that transcription read-through occurs very early after mitosis and that regulation of read-through is dependent on both cell type and cellular response.

The possible mechanisms underlying read-through transcription can include regulation of termination factors or improper 3’-processing of the RNA that could then interfere with termination. To investigate the latter possibility, we generated full-length PRO-seq (flPRO-seq, methods) data to examine the frequency of RNA cleavage at the poly-A site at 2hr post mitosis compared to asynchronous cells (Methods). We observe an increase in reads that span the poly-A site indicating that reduced RNA cleavage is, at least in part, responsible for transcription read-through after exit from mitosis (Figure S7F, 7J). This is consistent with a relatively decreased frequency of poly-A cleavage sites in regions of read-through transcription (Figure S7G). We also used the full-length data to examine splicing as previous work has indicated that the read-through transcripts undergo splicing at novel sites downstream of genes. Whereas we are able to observe normal splicing patterns within genes concomitant with read-through, we are unable to detect splicing of the read-through transcripts downstream of genes (Figure S7F, 7K, methods). The discrepancies could be due to differences in either the cellular response or mapping strategy used to detect splicing. These results demonstrate that dynamic regulation of transcription termination occurs during a brief window of time following exit from mitosis.

## DISCUSSION

### The role of transcription during mitosis

Using PRO-seq to capture active transcription, we have carefully dissected mitotic transcriptional activity in human cells. Incorporating multiple lines of evidence, we have identified a relatively low number (161) of genes that remain appreciably active during mitosis. The observation that these genes are strongly enriched for the canonical TATA box suggests that there is a requirement for strong core promoter elements in order to maintain transcription during mitosis. This is supported by the recent finding in mice that TATA binding protein (TBP) is maintained at select loci during mitosis (Teves, An, et al, 2018). Our mitotic-active genes are highly enriched in signaling and cell type-specific functions, including numerous transcription factors. These genes often overlap with the well-studied “early-response” genes, which have been implicated in a variety of cellular pathways (Vacca, Itoh, et al, 2018). While many of these genes are ubiquitously expressed, they demonstrate cell type-dependent transcription patterns specifically during mitosis. This suggests that expression of a select subset of genes during mitosis may play an important role in the maintenance of cell identity.

While the vast majority of genes are repressed at the level of transcription elongation in mitotic cells, we find that promoter architecture is largely maintained as found in other studies (Hsiung, Morrissey, et al, 2015; Liu, Pelham-Webb, et al, 2017; Teves, An, et al, 2016). Close inspection of our PRO-seq data shows that Pol II occupancy shifts downstream to a position adjacent to or overlapping the first nucleosome. This shift resembles that seen in NELF-depleted cells (Aoi, Smith, et al, 2020) and suggests the occurrence of improper polymerase-pausing and collision with the +1 nucleosome. The concomitant outward spreading of chromatin accessibility detected with ATAC-seq therefore indicates that RNA polymerase itself, may play an important role in the maintenance of promoter accessibility during mitosis. However, since the overall occupancy of Pol II at promoters is low during mitosis, yet strong enough to change the occupancy of the +1 nucleosome, we hypothesize that Pol II transcribes into the first nucleosome where it terminates by an unknown mechanism. A possible explanation could be that the hyper-phosphorylation of TFIID, TFIIH, and other transcription factors results in improper pause-regulation and failure to assemble a competent elongation complex. Termination of transcription leading up to and during mitosis is carried out by TTF2, which localizes to the nucleus specifically during mitosis (Jiang, Liu, et al, 2004). It contributes to the overall repression of transcription during mitosis and could also function in the polymerase turnover at promoters described here, although we have not ruled out other mechanisms involving the integrator complex and the canonical Pol II termination machinery.

RNA polymerase turnover at promoters and enhancers provides an attractive explanation for the maintenance of both chromatin accessibility and histone modifications during mitosis, while at the same time limiting full-length RNA production. Histone acetylation has been implicated in mitotic bookmarking numerous times, while also exhibiting a global reduction during mitosis (Behera, Stonestrom, et al, 2019; Javasky, Shamir, et al, 2018; Liu, Chen, et al, 2017; Liu, Pelham-Webb, et al, 2017; Pelham-Webb, Polyzos, et al, 2021). Combined with the observation that histone acetylation is dependent on transcription (Martin, Brind’Amour, et al, 2021; Wang et al., 2020 bioRxiv), our data suggest that this modification is maintained in conjunction with transcriptional activity and that both are also strong indicators of rapid reactivation following mitosis. While levels of trimethylation of H3K4 appear to remain unchanged during mitosis, it is unclear whether this modification has a direct role in transcription memory or reactivation kinetics following mitosis. In fact, as can be seen in the promoter composite profiles, nucleosomes containing H3K4me3 encroach upon the NDR during mitosis (Javasky, Shamir, et al, 2018; Liang, Woodfin, et al, 2015). This has also been reported in mitotic H2AZ profiles at both promoters and CTCF sites (Kelly, Miranda, et al, 2010; Oomen, Hansen, et al, 2019). Given the conflicting reports of accessibility and nucleosome deposition at promoters during mitosis, there appears to be a kinetic competition between mitotic chromatin condensation and the preservation of interphase NDRs. Our data suggests that basal activity of RNA polymerase at promoters and select enhancers maintains the local chromatin architecture of these elements and thereby acts as a mechanism of “cellular memory” that leads to rapid reactivation during exit from mitosis.

### Distinct profiles of genome reactivation following mitosis

Similar to previous results (Palozola, Donahue, et al, 2017), we find that the earliest genes to be reactivated following mitosis are enriched for general housekeeping functions and rebuilding the cell. However, we have also identified a subset of early genes whose expression is highest during the mitosis-G1 transition. These transiently expressed genes are enriched in both signaling and cell type-specific functions, providing support for an earlier hypothesis stating that genes important for cell identity would need to be reactivated first (Kadauke, Udugama, et al, 2012). Subsequent waves of gene reactivation are involved in cellular metabolism and gene expression at 3-4hrs post-release, followed by a secondary wave of cell type-specific genes at 6-8hrs. In addition to timing of activation, we also identify gene classes with distinct amplitudes of transcription activity following mitosis. Comparison of spike-in normalized PRO-seq signal throughout our time course revealed that a subset of housekeeping genes is reactivated with a “burst” of transcription while genes associated with metabolism and gene expression are reactivated in a more gradual manner. On the other hand, through measurements of both chromatin accessibility and transcriptional activity we have found that enhancer reactivation is delayed relative to genes. This likely reflects the overall weaker capacity of enhancer elements in driving transcription and coincides with recent reports demonstrating a lag in enhancer-promoter contact reformation (Abramo, Valton, et al, 2019; Pelham-Webb, Polyzos, et al, 2021; Zhang, Emerson, et al, 2019).

### Mitotic determinants of reactivation kinetics

We show that histone modifications at promoters represent general marks for reactivation, but are not fully predictive of kinetics. Alternatively, the best predictors of rapid gene reactivation are promoter accessibility and transcriptional activity during mitosis. The data suggest that promoter activity both during and immediately following mitosis are likely driven by overall promoter strength, rather than a single factor. In support of this model, early-activated genes also demonstrate the highest enrichment for basal transcription factors such as TBP and TAF1 binding. Importantly, direct binding of TBP has previously been associated with cell type-specific gene regulation (Goodrich, and Tjian, 2010). Given that P300 activity is important for activation of early G1 genes in iPSC cells (Pelham-Webb, Polyzos, et al, 2021), we suggest that cross-talk between acetylation, promoter strength and transcription can contribute to cell type specific differences in mitotically active promoters and enhancers. In support of this, the maintenance of high acetylation levels in mitotic mouse iPSCs is accompanied by a higher retention of transcription in these cells (Pelham-Webb, Polyzos, et al, 2021). Ultimately our results, along with recent studies, suggest that basal transcription during mitosis helps to maintain a privileged chromatin state at promoters and select enhancers, which then direct the initial rounds of post-mitotic transcription.

### Transcription phenotypes during mitosis-G1 transition

Promoter-proximal pausing is proposed to allow for dynamic and specific control of gene expression in changing environments. Although pause escape is an obligate step for nearly all genes (Guenther, Levine, et al, 2007; Jonkers, Kwak, and Lis, 2014; Peterlin, and Price, 2006), transcriptional changes are often dually regulated by both recruitment and initiation of Pol II as well (Hah, Danko, et al, 2011; Min, Waterfall, et al, 2011). Few examples exist where regulation of promoter-proximal pausing is the main mechanism responsible for broad changes in gene transcription (Mahat, Kwak, et al, 2016). Here, we find that Pol II rapidly transitions from a paused state into productive elongation at nearly all genes immediately following mitosis. We hypothesize that reduced transcription activity during mitosis results in a large pool of free P-TEFb that is rapidly recruited to promoters at the onset of gene reactivation, facilitating rapid release of Pol II into productive elongation. This state of hyper-release could be linked to post-mitotic bursting of transcription as seen here and in previous studies (Hsiung, Morrissey, et al, 2015). P-TEFb reassociation with gene bodies would then deplete the free pool, returning pausing to a rate-limiting step as seen in later G1 timepoints (post 4hrs). Overall, it appears that RNA polymerase activity during mitosis helps to maintain promoter architecture and primes genes for rapid reactivation.

The hyper-elongative state also appears to be linked to a global read-through transcription phenotype. We have demonstrated that widespread read-through transcription occurs specifically during early-G1 in multiple cell types, regardless of the method of cell cycle arrest. Read-through can continue hundreds of kb beyond the normal termination site and emanates from antisense promoter-associated transcripts and enhancers as well as genes. While this transcriptional phenotype resembles that of cells responding to various stresses such as osmotic and heat stress, or viral infection (Cardiello, Goodrich, and Kugel, 2018; Erickson, Sheridan, et al, 2018; Heinz, Texari, et al, 2018; Rutkowski, Erhard, et al, 2015; Vilborg, Sabath, et al, 2017), here we show that it occurs during global genome reactivation following mitosis. Given that this brief period of read-through transcription co-occurs with hyper-active pause-release, it is possible that the elongating transcription complex is modified in such a way during pause-release that alters the ability of factors involved in 3’-end formation and termination to associate. Consistent with this, creation of excess free and active P-TEFb through knockdown of the inhibitory 7sK RNA leads to a similar transcription phenotype in mice (Castelo-Branco, Amaral, et al, 2013; Guo, Li, and Price, 2013). However, viral induced transcription readthrough results from disruption of CPSF30 action by a viral protein and failure to cleave the RNA at the poly-A site. Thus, more studies are needed to determine the mechanism underlying transcription readthrough upon mitotic exit.

Although future work is required to elucidate a function of read-through transcription during mitosis, recent studies from other systems point to a role in regulating the chromatin architecture. RNA polymerase II and associated elongation factors are capable of dramatically altering the chromatin environment (Heinz, Texari, et al, 2018; Saunders, Core, and Lis, 2006). Pioneering rounds of transcription have been observed to assist in activation of genes presumably through disruption of chromatin architecture which gives regulatory factors access to the DNA (Adelman and Lis, 2012; Boettiger and Levine, 2009). We hypothesize that transcription read-through could provide such a pioneering activity for the genome after mitosis. The increased transcription footprint at this time could help to reset the chromatin architecture around genes through release of mitotic compaction. We also hypothesize that read-through transcription could have an effect on 3-dimensional genome architecture as the timing coincides with enhancer-promoter contact reformation (Abramo, Valton, et al, 2019; Pelham-Webb, Polyzos, et al, 2021; Zhang, Emerson, et al, 2019). Recent work supports a role for RNA polymerase in architectural resetting during exit from mitosis (Zhang et al., 2020 bioRxiv) and it has been demonstrated that RNA Pol II is able to alter the 3-dimensional chromatin environment by a number of mechanisms (Busslinger, Stocsits, et al, 2017; Heinz, Texari, et al, 2018; Hsieh, Cattoglio, et al, 2020). Thus, we believe that the initial rounds of transcription likely play a role in resetting both local and long-range genomic structure following mitosis. Overall, our data suggests that each step of transcription regulation is exploited during the mitosis-G1 transition, producing highly specialized and dynamic gene transcription profiles that likely contribute to the maintenance of cell identity and chromatin organization.

## Acknowledgements

We thank members of the Core Lab for helpful discussion and comments on the manuscript. We thank Bo Reese and The Center for Genome Innovation at the University of Connecticut for experimental discussion, library QC and sequencing. We also thank Wu He at the Center for Open Research Resources and Equipment at the University of Connecticut for assistance with performing flow cytometry. This work was supported by NIGMS grant R35GM128857 to LC.

## Author contributions

L.C. and L.A.W. designed and carried out the PRO-seq experiments. L.C. and C.K. produced the ATAC-seq data. L.C. and L.A.W. performed FACS analysis of synchronized cells. L.A.W. analyzed all new and publicly available data. L.M.W. performed microscopy imaging of synchronized cells. G.J.V. and L.A.W. created the full-length PRO-seq data. G.J.V. and P.N. contributed to data analyses and visualization. L.A.W. and L.C. wrote the manuscript. All Authors commented on the manuscript.

## REFERENCES

Abramo, K., Valton, A.L., Venev, S.V., Ozadam, H., Fox, A.N., and Dekker, J. (2019). A chromosome folding intermediate at the condensin-to-cohesin transition during telophase. Nat. Cell Biol. 11, 1393–1402.

Aoi, Y., Smith, E.R., Shah, A.P., Rendleman, E.J., Marshall, S.A., Woodfin, A.R., Chen, F.X., Shiekhattar, R., and Shilatifard, A. (2020). NELF Regulates a Promoter-Proximal Step Distinct from RNA Pol II Pause-Release. Mol. Cell 2, 261–274.e5.

Attwooll, C., Lazzerini Denchi, E., and Helin, K. (2004). The E2F family: specific functions and overlapping interests. EMBO J. 24, 4709–4716.

Bailey, T.L., Boden, M., Buske, F.A., Frith, M., Grant, C.E., Clementi, L., Ren, J., Li, W.W., and Noble, W.S. (2009). MEME SUITE: tools for motif discovery and searching. Nucleic Acids Res. Web Server issue, W202–8.

Behera, V., Stonestrom, A.J., Hamagami, N., Hsiung, C.C., Keller, C.A., Giardine, B., Sidoli, S., Yuan, Z.F., Bhanu, N.V., Werner, M.T. et al. (2019). Interrogating Histone Acetylation and BRD4 as Mitotic Bookmarks of Transcription. Cell. Rep. 2, 400–415.e5.

Bellier, S., Chastant, S., Adenot, P., Vincent, M., Renard, J.P., and Bensaude, O. (1997). Nuclear translocation and carboxyl-terminal domain phosphorylation of RNA polymerase II delineate the two phases of zygotic gene activation in mammalian embryos. EMBO J. 20, 6250–6262.

Bishop, A.C., Ubersax, J.A., Petsch, D.T., Matheos, D.P., Gray, N.S., Blethrow, J., Shimizu, E., Tsien, J.Z., Schultz, P.G., Rose, M.D. et al. (2000). A chemical switch for inhibitor-sensitive alleles of any protein kinase. Nature 6802, 395–401.

Buenrostro, J.D., Giresi, P.G., Zaba, L.C., Chang, H.Y., and Greenleaf, W.J. (2013). Transposition of native chromatin for fast and sensitive epigenomic profiling of open chromatin, DNA-binding proteins and nucleosome position. Nat. Methods 12, 1213–1218.

Busslinger, G.A., Stocsits, R.R., van der Lelij, P., Axelsson, E., Tedeschi, A., Galjart, N., and Peters, J.M. (2017). Cohesin is positioned in mammalian genomes by transcription, CTCF and Wapl. Nature 7651, 503–507.

Cardiello, J.F., Goodrich, J.A., and Kugel, J.F. (2018). Heat Shock Causes a Reversible Increase in RNA Polymerase II Occupancy Downstream of mRNA Genes, Consistent with a Global Loss in Transcriptional Termination. Mol. Cell. Biol. 18, 10.1128/MCB.00181-18. Print 2018 Sep 15.

Castelo-Branco, G., Amaral, P.P., Engstrom, P.G., Robson, S.C., Marques, S.C., Bertone, P., and Kouzarides, T. (2013). The non-coding snRNA 7SK controls transcriptional termination, poising, and bidirectionality in embryonic stem cells. Genome Biol. 9, R98-2013-14-9-r98.

Core, L.J., Martins, A.L., Danko, C.G., Waters, C.T., Siepel, A., and Lis, J.T. (2014). Analysis of nascent RNA identifies a unified architecture of initiation regions at mammalian promoters and enhancers. Nat. Genet. 12, 1311–1320.

Danko, C.G., Hyland, S.L., Core, L.J., Martins, A.L., Waters, C.T., Lee, H.W., Cheung, V.G., Kraus, W.L., Lis, J.T., and Siepel, A. (2015). Identification of active transcriptional regulatory elements from GRO-seq data. Nat. Methods 5, 433–438.

Erickson, B., Sheridan, R.M., Cortazar, M., and Bentley, D.L. (2018). Dynamic turnover of paused Pol II complexes at human promoters. Genes Dev. 17-18, 1215–1225.

Gazit, B., Cedar, H., Lerer, I., and Voss, R. (1982). Active genes are sensitive to deoxyribonuclease I during metaphase. Science 4560, 648–650.

Goodrich, J.A., and Tjian, R. (2010). Unexpected roles for core promoter recognition factors in cell-type-specific transcription and gene regulation. Nat. Rev. Genet. 8, 549–558.

Gravells, P., Tomita, K., Booth, A., Poznansky, J., and Porter, A.C. (2013). Chemical genetic analyses of quantitative changes in Cdk1 activity during the human cell cycle. Hum. Mol. Genet. 14, 2842–2851.

Groudine, M., and Weintraub, H. (1982). Propagation of globin DNAase I-hypersensitive sites in absence of factors required for induction: a possible mechanism for determination. Cell 1, 131–139.

Guenther, M.G., Levine, S.S., Boyer, L.A., Jaenisch, R., and Young, R.A. (2007). A chromatin landmark and transcription initiation at most promoters in human cells. Cell 1, 77–88.

Guo, J., Li, T., and Price, D.H. (2013). Runaway transcription. Genome Biol. 9, 133–2013-14-9-133.

Hah, N., Danko, C.G., Core, L., Waterfall, J.J., Siepel, A., Lis, J.T., and Kraus, W.L. (2011). A rapid, extensive, and transient transcriptional response to estrogen signaling in breast cancer cells. Cell 4, 622–634.

Heinz, S., Texari, L., Hayes, M.G.B., Urbanowski, M., Chang, M.W., Givarkes, N., Rialdi, A., White, K.M., Albrecht, R.A., Pache, L. et al. (2018). Transcription Elongation Can Affect Genome 3D Structure. Cell 6, 1522–1536.e22.

Henriques, T., Scruggs, B.S., Inouye, M.O., Muse, G.W., Williams, L.H., Burkholder, A.B., Lavender, C.A., Fargo, D.C., and Adelman, K. (2018). Widespread transcriptional pausing and elongation control at enhancers. Genes Dev. 1, 26–41.

Hershkovitz, M., and Riggs, A.D. (1997). Ligation-mediated PCR for chromatin-structure analysis of interphase and metaphase chromatin. Methods 2, 253–263.

Hsieh, T.S., Cattoglio, C., Slobodyanyuk, E., Hansen, A.S., Rando, O.J., Tjian, R., and Darzacq, X. (2020). Resolving the 3D Landscape of Transcription-Linked Mammalian Chromatin Folding. Mol. Cell 3, 539–553.e8.

Hsiung, C.C., Bartman, C.R., Huang, P., Ginart, P., Stonestrom, A.J., Keller, C.A., Face, C., Jahn, K.S., Evans, P., Sankaranarayanan, L. et al. (2016). A hyperactive transcriptional state marks genome reactivation at the mitosis-G1 transition. Genes Dev. 12, 1423–1439.

Hsiung, C.C., Morrissey, C.S., Udugama, M., Frank, C.L., Keller, C.A., Baek, S., Giardine, B., Crawford, G.E., Sung, M.H., Hardison, R.C., and Blobel, G.A. (2015). Genome accessibility is widely preserved and locally modulated during mitosis. Genome Res. 2, 213–225.

Javasky, E., Shamir, I., Gandhi, S., Egri, S., Sandler, O., Rothbart, S.B., Kaplan, N., Jaffe, J.D., Goren, A., and Simon, I. (2018). Study of mitotic chromatin supports a model of bookmarking by histone modifications and reveals nucleosome deposition patterns. Genome Res. 10, 1455–1466.

Jiang, Y., Liu, M., Spencer, C.A., and Price, D.H. (2004). Involvement of transcription termination factor 2 in mitotic repression of transcription elongation. Mol. Cell 3, 375–385.

Jonkers, I., Kwak, H., and Lis, J.T. (2014). Genome-wide dynamics of Pol II elongation and its interplay with promoter proximal pausing, chromatin, and exons. Elife e02407.

Kadauke, S., Udugama, M.I., Pawlicki, J.M., Achtman, J.C., Jain, D.P., Cheng, Y., Hardison, R.C., and Blobel, G.A. (2012). Tissue-specific mitotic bookmarking by hematopoietic transcription factor GATA1. Cell 4, 725–737.

Kang, H., Shokhirev, M.N., Xu, Z., Chandran, S., Dixon, J.R., and Hetzer, M.W. (2020). Dynamic regulation of histone modifications and long-range chromosomal interactions during postmitotic transcriptional reactivation. Genes Dev. 13-14, 913–930.

Kelly, T.K., Miranda, T.B., Liang, G., Berman, B.P., Lin, J.C., Tanay, A., and Jones, P.A. (2010). H2A.Z maintenance during mitosis reveals nucleosome shifting on mitotically silenced genes. Mol. Cell 6, 901–911.

Kent, W.J., Sugnet, C.W., Furey, T.S., Roskin, K.M., Pringle, T.H., Zahler, A.M., and Haussler, D. (2002). The human genome browser at UCSC. Genome Res. 6, 996–1006.

Kfir, N., Lev-Maor, G., Glaich, O., Alajem, A., Datta, A., Sze, S.K., Meshorer, E., and Ast, G. (2015). SF3B1 association with chromatin determines splicing outcomes. Cell. Rep. 4, 618–629.

Kim, D., Langmead, B., and Salzberg, S.L. (2015). HISAT: a fast spliced aligner with low memory requirements. Nat. Methods 4, 357–360.

Kim, E., Du, L., Bregman, D.B., and Warren, S.L. (1997). Splicing factors associate with hyperphosphorylated RNA polymerase II in the absence of pre-mRNA. J. Cell Biol. 1, 19–28.

Kuo, M.T. (1982). Analysis of DNA attached to the chromosome scaffold. J. Cell Biol. 2, 278–284.

Langmead, B., Trapnell, C., Pop, M., and Salzberg, S.L. (2009). Ultrafast and memory-efficient alignment of short DNA sequences to the human genome. Genome Biol. 3, R25-2009-10-3-r25. Epub 2009 Mar 4.

Leresche, A., Wolf, V.J., and Gottesfeld, J.M. (1996). Repression of RNA polymerase II and III transcription during M phase of the cell cycle. Exp. Cell Res. 2, 282–288.

Li, H., Handsaker, B., Wysoker, A., Fennell, T., Ruan, J., Homer, N., Marth, G., Abecasis, G., Durbin, R., and 1000 Genome Project Data Processing Subgroup. (2009). The Sequence Alignment/Map format and SAMtools. Bioinformatics 16, 2078–2079.

Liang, K., Woodfin, A.R., Slaughter, B.D., Unruh, J.R., Box, A.C., Rickels, R.A., Gao, X., Haug, J.S., Jaspersen, S.L., and Shilatifard, A. (2015). Mitotic Transcriptional Activation: Clearance of Actively Engaged Pol II via Transcriptional Elongation Control in Mitosis. Mol. Cell 3, 435–445.

Liu, Y., Chen, S., Wang, S., Soares, F., Fischer, M., Meng, F., Du, Z., Lin, C., Meyer, C., DeCaprio, J.A. et al. (2017). Transcriptional landscape of the human cell cycle. Proc. Natl. Acad. Sci. U. S. A. 13, 3473–3478.

Liu, Y., Pelham-Webb, B., Di Giammartino, D.C., Li, J., Kim, D., Kita, K., Saiz, N., Garg, V., Doane, A., Giannakakou, P. et al. (2017). Widespread Mitotic Bookmarking by Histone Marks and Transcription Factors in Pluripotent Stem Cells. Cell. Rep. 7, 1283–1293.

Mahat, D.B., Kwak, H., Booth, G.T., Jonkers, I.H., Danko, C.G., Patel, R.K., Waters, C.T., Munson, K., Core, L.J., and Lis, J.T. (2016). Base-pair-resolution genome-wide mapping of active RNA polymerases using precision nuclear run-on (PRO-seq). Nat. Protoc. 8, 1455–1476.

Martin, B.J.E., Brind’Amour, J., Kuzmin, A., Jensen, K.N., Liu, Z.C., Lorincz, M., and Howe, L.J. (2021). Transcription shapes genome-wide histone acetylation patterns. Nat. Commun. 1, 210–020-20543-z.

Martinez-Balbas, M.A., Dey, A., Rabindran, S.K., Ozato, K., and Wu, C. (1995). Displacement of sequence-specific transcription factors from mitotic chromatin. Cell 1, 29–38.

McLean, C.Y., Bristor, D., Hiller, M., Clarke, S.L., Schaar, B.T., Lowe, C.B., Wenger, A.M., and Bejerano, G. (2010). GREAT improves functional interpretation of cis-regulatory regions. Nat. Biotechnol. 5, 495–501.

Michelotti, E.F., Sanford, S., and Levens, D. (1997). Marking of active genes on mitotic chromosomes. Nature 6645, 895–899.

Mikhaylichenko, O., Bondarenko, V., Harnett, D., Schor, I.E., Males, M., Viales, R.R., and Furlong, E.E.M. (2018). The degree of enhancer or promoter activity is reflected by the levels and directionality of eRNA transcription. Genes Dev. 1, 42–57.

Min, I.M., Waterfall, J.J., Core, L.J., Munroe, R.J., Schimenti, J., and Lis, J.T. (2011). Regulating RNA polymerase pausing and transcription elongation in embryonic stem cells. Genes Dev. 7, 742–754.

Oomen, M.E., Hansen, A.S., Liu, Y., Darzacq, X., and Dekker, J. (2019). CTCF sites display cell cycle-dependent dynamics in factor binding and nucleosome positioning. Genome Res. 2, 236–249.

Owens, N., Papadopoulou, T., Festuccia, N., Tachtsidi, A., Gonzalez, I., Dubois, A., Vandormael-Pournin, S., Nora, E.P., Bruneau, B.G., Cohen-Tannoudji, M., and Navarro, P. (2019). CTCF confers local nucleosome resiliency after DNA replication and during mitosis. Elife 10.7554/eLife.47898.

Palozola, K.C., Donahue, G., Liu, H., Grant, G.R., Becker, J.S., Cote, A., Yu, H., Raj, A., and Zaret, K.S. (2017). Mitotic transcription and waves of gene reactivation during mitotic exit. Science 6359, 119–122.

Palozola, K.C., Lerner, J., and Zaret, K.S. (2019). A changing paradigm of transcriptional memory propagation through mitosis. Nat. Rev. Mol. Cell Biol. 1, 55–64.

Palozola, K.C., Liu, H., Nicetto, D., and Zaret, K.S. (2017). Low-Level, Global Transcription during Mitosis and Dynamic Gene Reactivation during Mitotic Exit. Cold Spring Harb. Symp. Quant. Biol. 197–205.

Parsons, G.G., and Spencer, C.A. (1997). Mitotic repression of RNA polymerase II transcription is accompanied by release of transcription elongation complexes. Mol. Cell. Biol. 10, 5791–5802.

Pelham-Webb, B., Murphy, D., and Apostolou, E. (2020). Dynamic 3D Chromatin Reorganization during Establishment and Maintenance of Pluripotency. Stem Cell. Reports 6, 1176–1195.

Pelham-Webb, B., Polyzos, A., Wojenski, L., Kloetgen, A., Li, J., Di Giammartino, D.C., Sakellaropoulos, T., Tsirigos, A., Core, L., and Apostolou, E. (2021). H3K27ac bookmarking promotes rapid post-mitotic activation of the pluripotent stem cell program without impacting 3D chromatin reorganization. Mol. Cell 8, 1732–1748.e8.

Peterlin, B.M., and Price, D.H. (2006). Controlling the elongation phase of transcription with P-TEFb. Mol. Cell 3, 297–305.

Prescott, D.M., and Bender, M.A. (1962). Synthesis of RNA and protein during mitosis in mammalian tissue culture cells. Exp. Cell Res. 260–268.

Quinlan, A.R., and Hall, I.M. (2010). BEDTools: a flexible suite of utilities for comparing genomic features. Bioinformatics 6, 841–842.

Raccaud, M., and Suter, D.M. (2018). Transcription factor retention on mitotic chromosomes: regulatory mechanisms and impact on cell fate decisions. FEBS Lett. 6, 878–887.

Ramirez, F., Ryan, D.P., Gruning, B., Bhardwaj, V., Kilpert, F., Richter, A.S., Heyne, S., Dundar, F., and Manke, T. (2016). deepTools2: a next generation web server for deep-sequencing data analysis. Nucleic Acids Res. W1, W160–5.

Robinson, M.D., McCarthy, D.J., and Smyth, G.K. (2010). edgeR: a Bioconductor package for differential expression analysis of digital gene expression data. Bioinformatics 1, 139–140.

Rothe, M., Pehl, M., Taubert, H., and Jackle, H. (1992). Loss of gene function through rapid mitotic cycles in the Drosophila embryo. Nature 6391, 156–159.

Rutkowski, A.J., Erhard, F., L’Hernault, A., Bonfert, T., Schilhabel, M., Crump, C., Rosenstiel, P., Efstathiou, S., Zimmer, R., Friedel, C.C., and Dolken, L. (2015). Widespread disruption of host transcription termination in HSV-1 infection. Nat. Commun. 7126.

Saunders, A., Core, L.J., and Lis, J.T. (2006). Breaking barriers to transcription elongation. Nat. Rev. Mol. Cell Biol. 8, 557–567.

Schep, A.N., Buenrostro, J.D., Denny, S.K., Schwartz, K., Sherlock, G., and Greenleaf, W.J. (2015). Structured nucleosome fingerprints enable high-resolution mapping of chromatin architecture within regulatory regions. Genome Res. 11, 1757–1770.

Segil, N., Guermah, M., Hoffmann, A., Roeder, R.G., and Heintz, N. (1996). Mitotic regulation of TFIID: inhibition of activator-dependent transcription and changes in subcellular localization. Genes Dev. 19, 2389–2400.

Segil, N., Roberts, S.B., and Heintz, N. (1991). Mitotic phosphorylation of the Oct-1 homeodomain and regulation of Oct-1 DNA binding activity. Science 5039, 1814–1816.

Sheffield, N.C., and Bock, C. (2016). LOLA: enrichment analysis for genomic region sets and regulatory elements in R and Bioconductor. Bioinformatics 4, 587–589.

Shermoen, A.W., and O’Farrell, P.H. (1991). Progression of the cell cycle through mitosis leads to abortion of nascent transcripts. Cell 2, 303–310.

Taylor, J.H. (1960). Nucleic acid synthesis in relation to the cell division cycle. Ann. N. Y. Acad. Sci. 409–421.

Teves, S.S., An, L., Bhargava-Shah, A., Xie, L., Darzacq, X., and Tjian, R. (2018). A stable mode of bookmarking by TBP recruits RNA polymerase II to mitotic chromosomes. Elife 10.7554/eLife.35621.

Teves, S.S., An, L., Hansen, A.S., Xie, L., Darzacq, X., and Tjian, R. (2016). A dynamic mode of mitotic bookmarking by transcription factors. Elife 10.7554/eLife.22280.

Tippens, N.D., Liang, J., Leung, A.K., Wierbowski, S.D., Ozer, A., Booth, J.G., Lis, J.T., and Yu, H. (2020). Transcription imparts architecture, function and logic to enhancer units. Nat. Genet. 10, 1067–1075.

Vacca, A., Itoh, M., Kawaji, H., Arner, E., Lassmann, T., Daub, C.O., Carninci, P., Forrest, A.R.R., Hayashizaki, Y., FANTOM Consortium, Aitken, S., and Semple, C.A. (2018). Conserved temporal ordering of promoter activation implicates common mechanisms governing the immediate early response across cell types and stimuli. Open Biol. 8, 10.1098/rsob.180011.

Vilborg, A., Passarelli, M.C., Yario, T.A., Tycowski, K.T., and Steitz, J.A. (2015). Widespread Inducible Transcription Downstream of Human Genes. Mol. Cell 3, 449–461.

Vilborg, A., Sabath, N., Wiesel, Y., Nathans, J., Levy-Adam, F., Yario, T.A., Steitz, J.A., and Shalgi, R. (2017). Comparative analysis reveals genomic features of stress-induced transcriptional readthrough. Proc. Natl. Acad. Sci. U. S. A. 40, E8362–E8371.

Warren, S.L., Landolfi, A.S., Curtis, C., and Morrow, J.S. (1992). Cytostellin: a novel, highly conserved protein that undergoes continuous redistribution during the cell cycle. J. Cell. Sci. Pt 2, 381–388.

White, R.J., Gottlieb, T.M., Downes, C.S., and Jackson, S.P. (1995). Mitotic regulation of a TATA-binding-protein-containing complex. Mol. Cell. Biol. 4, 1983–1992.

Whitfield, M.L., Sherlock, G., Saldanha, A.J., Murray, J.I., Ball, C.A., Alexander, K.E., Matese, J.C., Perou, C.M., Hurt, M.M., Brown, P.O., and Botstein, D. (2002). Identification of genes periodically expressed in the human cell cycle and their expression in tumors. Mol. Biol. Cell 6, 1977–2000.

Wissink, E.M., Vihervaara, A., Tippens, N.D., and Lis, J.T. (2019). Nascent RNA analyses: tracking transcription and its regulation. Nat. Rev. Genet. 12, 705–723.

Yu, G., Wang, L.G., Han, Y., and He, Q.Y. (2012). clusterProfiler: an R package for comparing biological themes among gene clusters. OMICS 5, 284–287.

Zang, C., Schones, D.E., Zeng, C., Cui, K., Zhao, K., and Peng, W. (2009). A clustering approach for identification of enriched domains from histone modification ChIP-Seq data. Bioinformatics 15, 1952–1958.

Zawel, L., Lu, H., Cisek, L.J., Corden, J.L., and Reinberg, D. (1993). The cycling of RNA polymerase II during transcription. Cold Spring Harb. Symp. Quant. Biol. 187–198.

Zhang, H., Emerson, D.J., Gilgenast, T.G., Titus, K.R., Lan, Y., Huang, P., Zhang, D., Wang, H., Keller, C.A., Giardine, B. et al. (2019). Chromatin structure dynamics during the mitosis-to-G1 phase transition. Nature 7785, 158–162.

Zhang, Y., Liu, T., Meyer, C.A., Eeckhoute, J., Johnson, D.S., Bernstein, B.E., Nusbaum, C., Myers, R.M., Brown, M., Li, W., and Liu, X.S. (2008). Model-based analysis of ChIP-Seq (MACS). Genome Biol. 9, R137-2008-9-9-r137. Epub 2008 Sep 17.

