## Supplemental figures and methods for "Multiple modes of regulation control dynamic transcription patterns during the mitosis-G1 transition"

**<u>Table of contents:</u>**

**Supplementary figures S1-S7**
**Methods**
**Quantification and Statistical Analysis**
**Supplementary Table 1**

### Figure S1

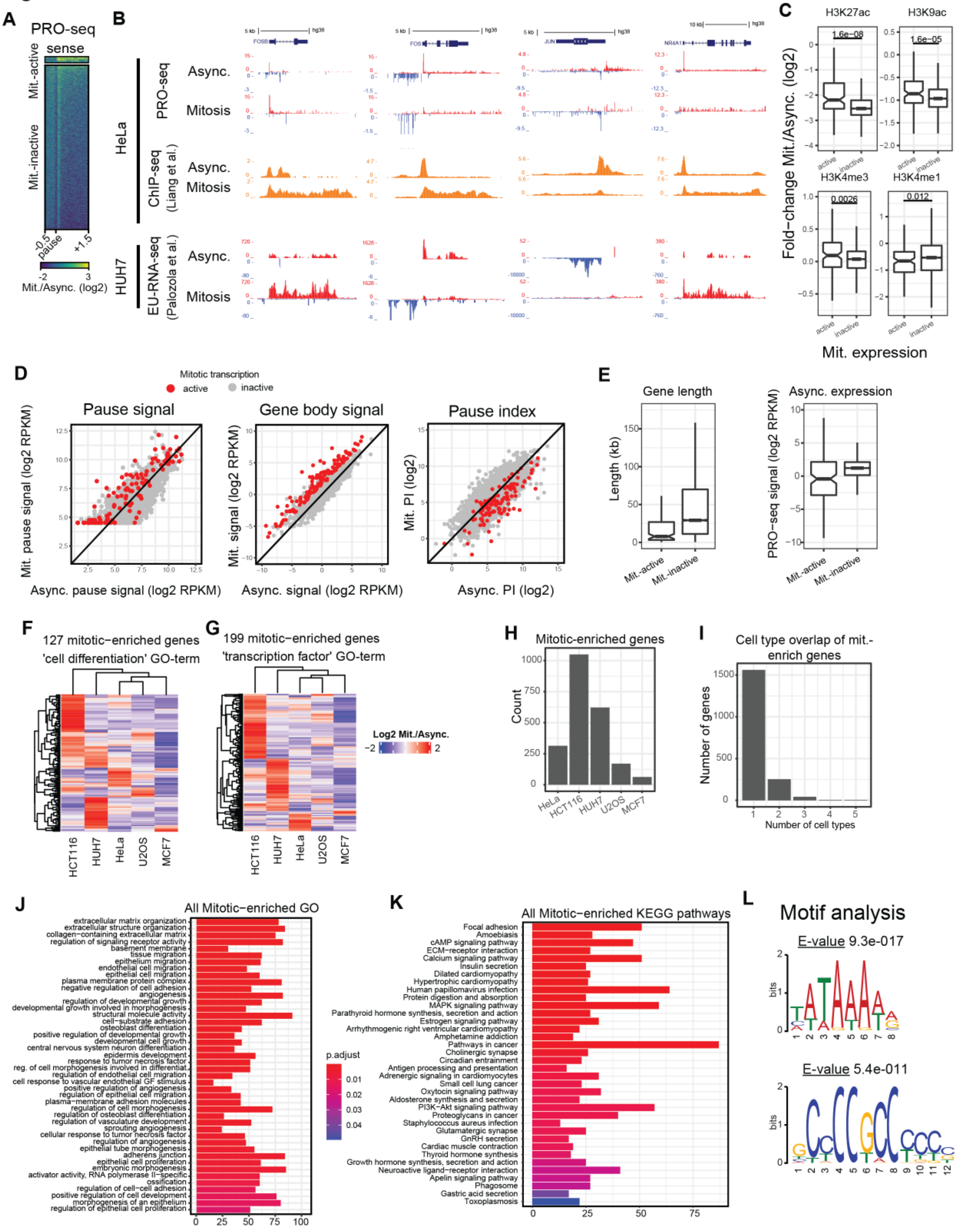
A
PRO-seq
sense
Mit.-active
Mit.-inactive
-0.5
pause
+1.5
-2
3
Mit./Async. (log2)
B
HeLa
PRO-seq
Async.
Mitosis
ChIP-seq (Liang et al.)
Async.
Mitosis
HUH7
EU-RNA-seq (Palozola et al.)
Async.
Mitosis
FOSB
FOS
JUN
NR4A1
hg38
C
H3K27ac
H3K9ac
H3K4me3
H3K4me1
1.6e−08
1.6e−05
0.0026
0.012
active
inactive
Fold−change Mit./Async. (log2)
Mit. expression
D
Mitotic transcription
active
inactive
Pause signal
Gene body signal
Pause index
Mit. pause signal (log2 RPKM)
Async. pause signal (log2 RPKM)
Mit. signal (log2 RPKM)
Async. signal (log2 RPKM)
Mit. PI (log2)
Async. PI (log2)
E
Gene length
Length (kb)
Async. expression
PRO−seq signal (log2 RPKM)
Mit.-active
Mit.-inactive
F
127 mitotic−enriched genes
'cell differentiation' GO-term
G
199 mitotic−enriched genes
'transcription factor' GO-term
Log2 Mit./Async.
−2
2
HCT116
HUH7
HeLa
U2OS
MCF7
H
Mitotic-enriched genes
Count
HeLa
HCT116
HUH7
U2OS
MCF7
I
Cell type overlap of mit.-enrich genes
Number of genes
Number of cell types
J
All Mitotic−enriched GO
extracellular matrix organization
extracellular structure organization
collagen−containing extracellular matrix
regulation of signaling receptor activity
basement membrane
tissue migration
epithelium migration
endothelial cell migration
epithelial cell migration
plasma membrane protein complex
negative regulation of cell adhesion
angiogenesis
regulation of developmental growth
developmental growth involved in morphogenesis
structural molecule activity
cell−substrate adhesion
osteoblast differentiation
positive regulation of developmental growth
developmental cell growth
central nervous system neuron differentiation
epidermis development
response to tumor necrosis factor
reg. of cell morphogenesis involved in differentiat.
regulation of endothelial cell migration
cell response to vascular endothelial GF stimulus
positive regulation of angiogenesis
regulation of epithelial cell migration
plasma−membrane adhesion molecules
regulation of cell morphogenesis
regulation of osteoblast differentiation
regulation of vasculature development
sprouting angiogenesis
cellular response to tumor necrosis factor
regulation of angiogenesis
epithelial tube morphogenesis
adherens junction
epithelial cell proliferation
embryonic morphogenesis
activator activity, RNA polymerase II−specific
ossification
regulation of cell−cell adhesion
positive regulation of cell development
morphogenesis of an epithelium
regulation of epithelial cell proliferation
p.adjust
K
All Mitotic−enriched KEGG pathways
Focal adhesion
Amoebiasis
cAMP signaling pathway
ECM−receptor interaction
Calcium signaling pathway
Insulin secretion
Dilated cardiomyopathy
Hypertrophic cardiomyopathy
Human papillomavirus infection
Protein digestion and absorption
MAPK signaling pathway
Parathyroid hormone synthesis, secretion and action
Estrogen signaling pathway
Arrhythmogenic right ventricular cardiomyopathy
Amphetamine addiction
Pathways in cancer
Cholinergic synapse
Circadian entrainment
Antigen processing and presentation
Adrenergic signaling in cardiomyocytes
Small cell lung cancer
Oxytocin signaling pathway
Aldosterone synthesis and secretion
PI3K−Akt signaling pathway
Proteoglycans in cancer
Staphylococcus aureus infection
Glutamatergic synapse
GnRH secretion
Cardiac muscle contraction
Thyroid hormone synthesis
Growth hormone synthesis, secretion and action
Neuroactive ligand−receptor interaction
Apelin signaling pathway
Phagosome
Gastric acid secretion
Toxoplasmosis
L
Motif analysis
E-value 9.3e-017
E-value 5.4e-011
bits

**Figure S1 (related to Figure 1)**

**(A)** Heatmap of log2 Mitotic/Async. sense PRO-seq signal in mitotic-active and inactive genes. **(B)** Genome browser tracks of asynchronous and mitotic datasets comparing transcription signal in HeLa cells based on PRO-seq and Pol II ChIP-seq and in HUH7 cells based on EU-RNA-seq. **(C)** Box plots of Mitotic/Async. ChIP-seq signal (log2) in promoters of mitotic-active and inactive genes. **(D)** Scatterplots of log2 pause signal, gene signal, and pausing index genome-wide in mitotic and asynchronous HeLa cells. Genes active in mitosis are in red. **(E)** Box plots of gene length and asynchronous transcription level of mitotic-active and inactive genes. **(F)** Heatmaps of Mitotic/Async. differential gene expression analysis in five different cell types showing genes associated with the GO term "cell differentiation". **(G)** Heatmaps as in G showing genes associated with the GO term "transcription factor". **(H)** Bar plot showing the number of mitotic-enriched genes for each cell type. **(I)** Bar plot summarizing the number of cell types in which genes are found to be mitotic-enriched. **(J)** Gene ontology analysis of mitotic-enriched genes across all five cell types. **(K)** KEGG pathway analysis of mitotic-enriched genes across all five cell types. **(L)** Motif search results in mitotic-active gene promoters.

**Figure S2**

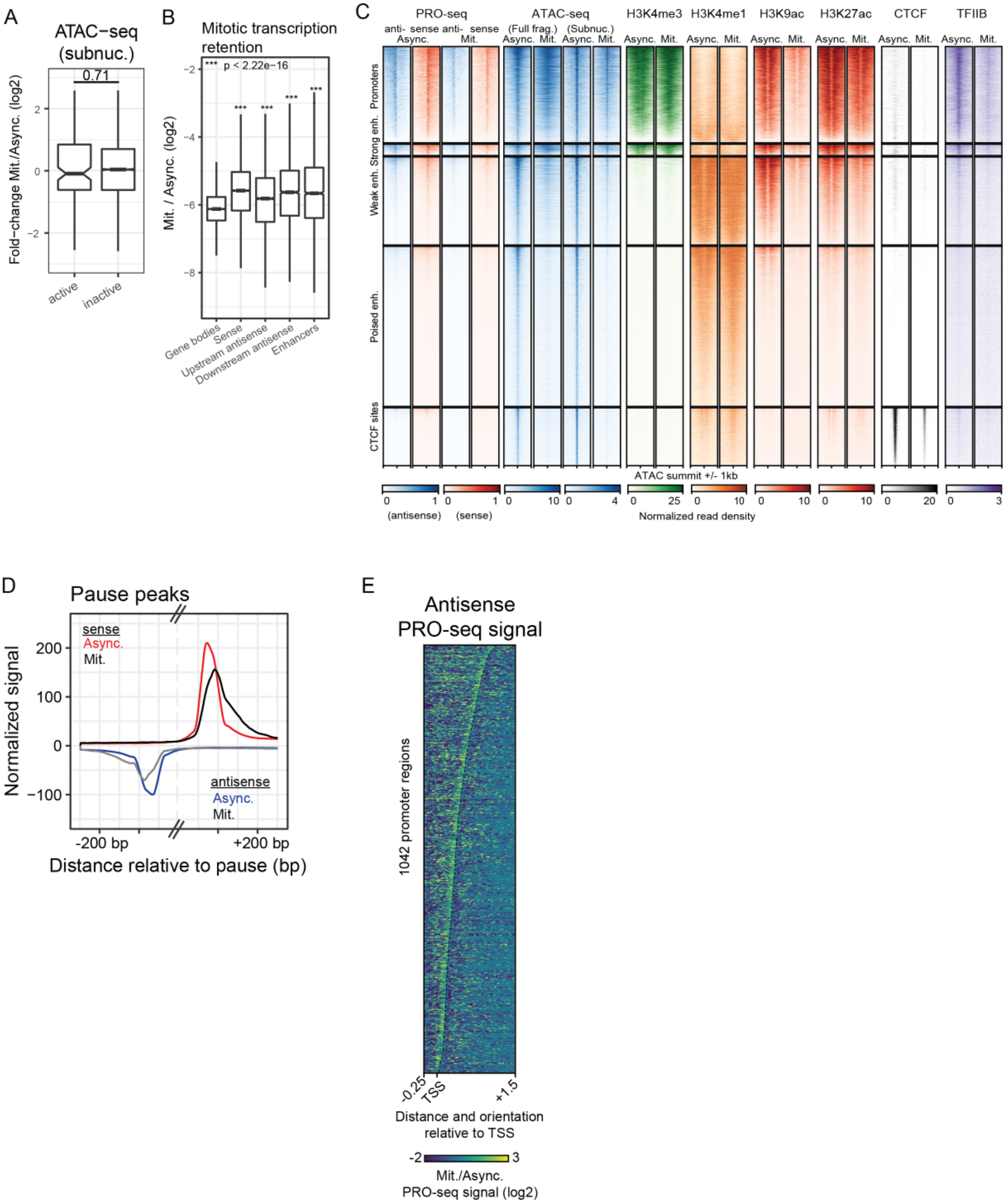
A
ATAC−seq (subnuc.)
0.71
Fold−change Mit./Async. (log2)
active
inactive
B
Mitotic transcription retention
*** p < 2.22e−16
Mit. / Async. (log2)
Gene bodies
Sense
Upstream antisense
Downstream antisense
Enhancers
C
PRO-seq
ATAC-seq
H3K4me3
H3K4me1
H3K9ac
H3K27ac
CTCF
TFIIB
Promoters
Strong enh.
Weak enh.
Poised enh.
CTCF sites
ATAC summit +/- 1kb
(antisense)
(sense)
Normalized read density
D
Pause peaks
sense
Async.
Mit.
antisense
Async.
Mit.
Normalized signal
-200 bp
+200 bp
Distance relative to pause (bp)
E
Antisense PRO-seq signal
1042 promoter regions
Distance and orientation relative to TSS
Mit./Async. PRO-seq signal (log2)

**Figure S2 (related to Figure 2)**

**(A)** Boxplots of log2 Mitotic/Async. fold-change of subnucleosomal ATAC-seq signal in promoters of genes active and inactive during mitosis. **(B)** Boxplots of log2 Mitotic/Async. fold-change of PRO-seq signal in transcribed genomic regions. **(C)** Heatmap of normalized mitotic and async. signal at genomic elements. **(D)** Average profiles of normalized PRO-seq signal relative to sense and antisense pause peaks. Distances between sense and antisense pause peaks are scaled to 100bp. **(E)** Heatmap of antisense PRO-seq signal at 1,042 genes with a daTSS in the first 1.5kb downstream of the sense pause peak. Rows are sorted by the distance between sense and antisense pause peaks.

#### Figure S3

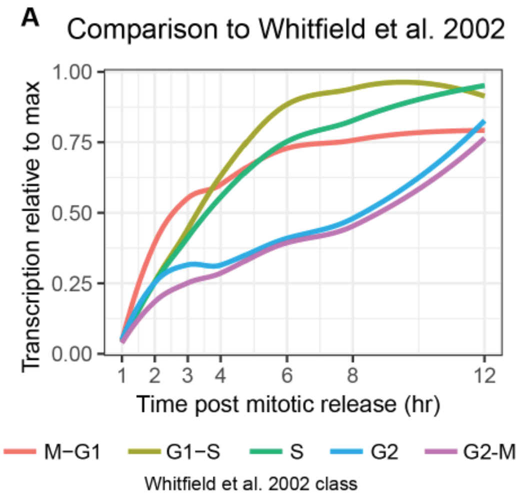


**Figure S3 (Related to figure 3)**

**(A)** Line plots of PRO-seq signal of gene kinetic classes from Whitfield et al., 2002.

#### Figure S4

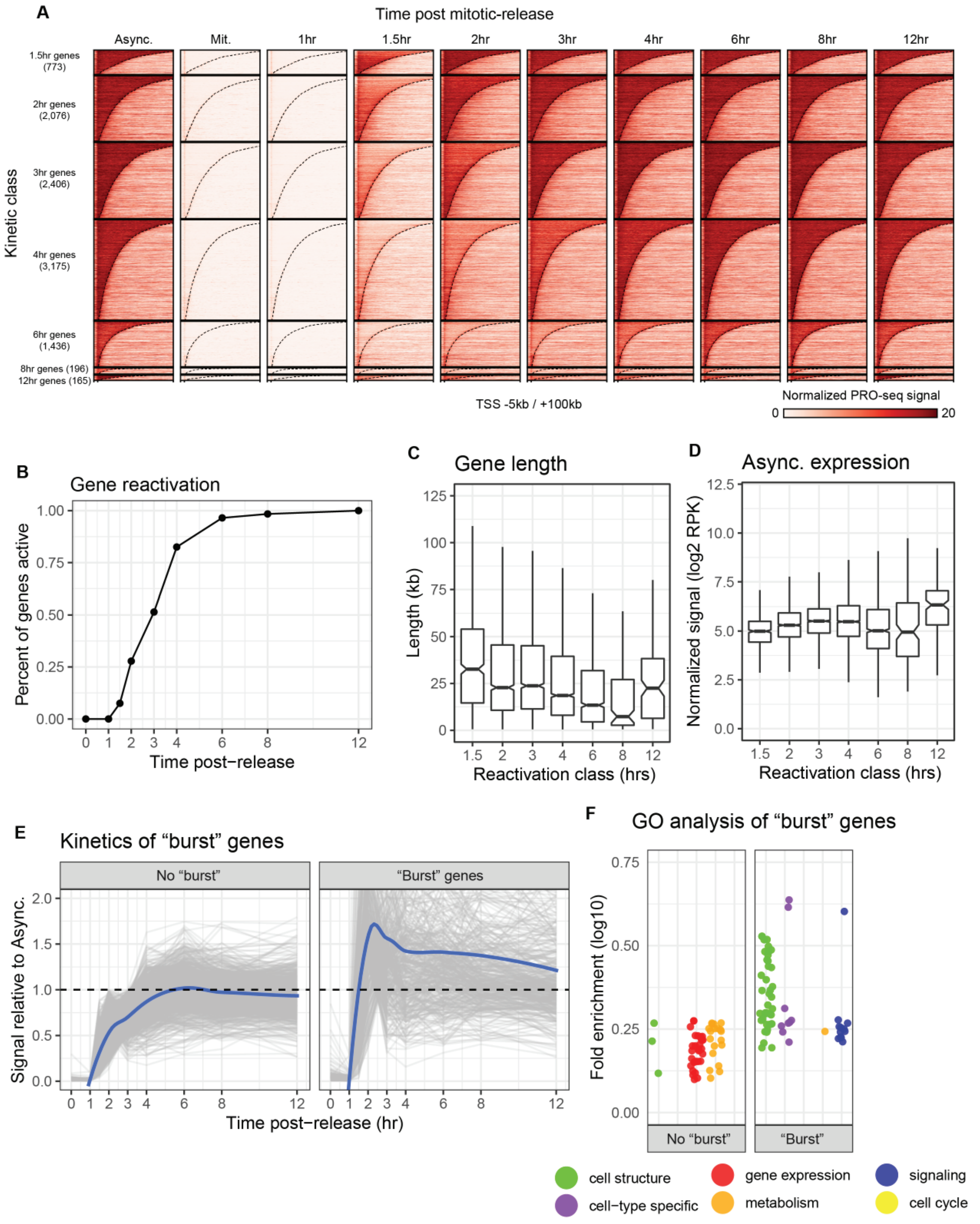
A
Time post mitotic-release
Async.
Mit.
1hr
1.5hr
2hr
3hr
4hr
6hr
8hr
12hr
Kinetic class
1.5hr genes (773)
2hr genes (2,076)
3hr genes (2,406)
4hr genes (3,175)
6hr genes (1,436)
8hr genes (196)
12hr genes (165)
TSS -5kb / +100kb
Normalized PRO-seq signal
0
20
B
Gene reactivation
Percent of genes active
Time post−release
C
Gene length
Length (kb)
Reactivation class (hrs)
D
Async. expression
Normalized signal (log2 RPK)
Reactivation class (hrs)
E
Kinetics of "burst" genes
No "burst"
"Burst" genes
Signal relative to Async.
Time post−release (hr)
F
GO analysis of "burst" genes
Fold enrichment (log10)
No "burst"
"Burst"
cell structure
cell−type specific
gene expression
metabolism
signaling
cell cycle

**Figure S4 (related to figure 4)**

**(A)** Heatmap of spike-in normalized PRO-seq signal in gene kinetic classes. Genes are order by length. **(B)** Cumulative distribution plot showing the fraction of genes reactivated at each timepoint following release from mitosis. **(C)** Boxplots of the gene length distribution for each kinetic class. **(D)** Boxplots of transcription level of genes in each kinetic class. **(E)** Line plots showing the kinetics of early genes relative to asynchronous expression levels. Genes have been grouped into indicated classes (methods). **(F)** Gene ontology analysis of early kinetic classes further grouped by signal relative to asynchronous cells.

#### Figure S5

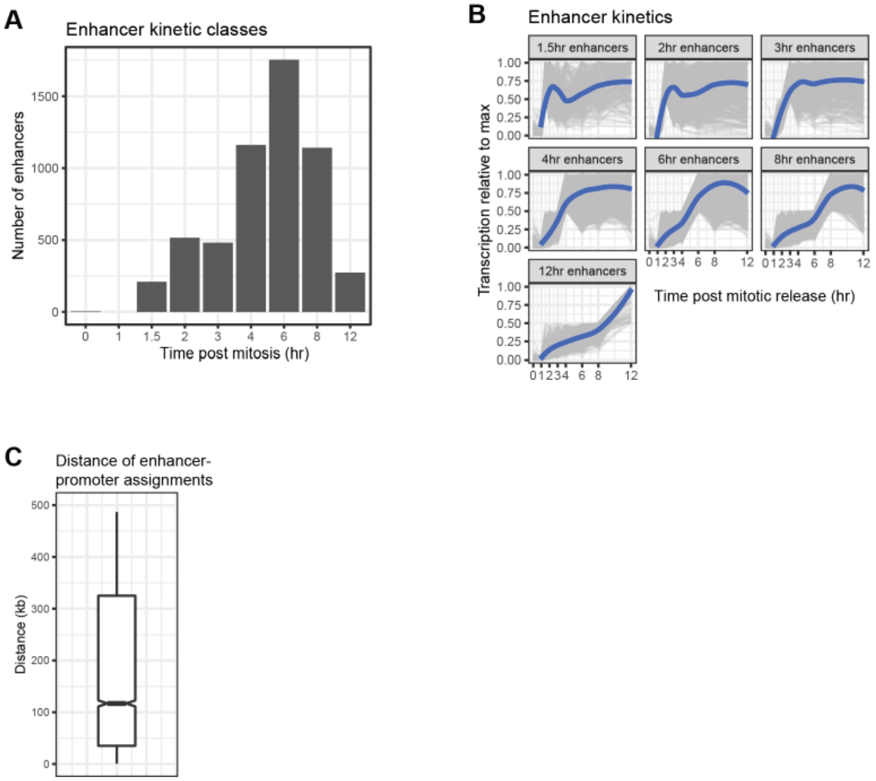


**Figure S5 (related to figure 5)**

**(A)** Summary bar plot of the number of enhancers in kinetic reactivation classes. **(B)** Line plots of enhancer kinetics during the 12hrs after release from mitosis for each kinetic class. A summary line (blue) was generated for each kinetic class by fitting a loess curve of the 1-12hr timepoints. Y-axis is spike-in normalized signal relative to maximum transcription level throughout the time course. **(C)** Boxplot of enhancer-promoter assignment distances.

#### Figure S6

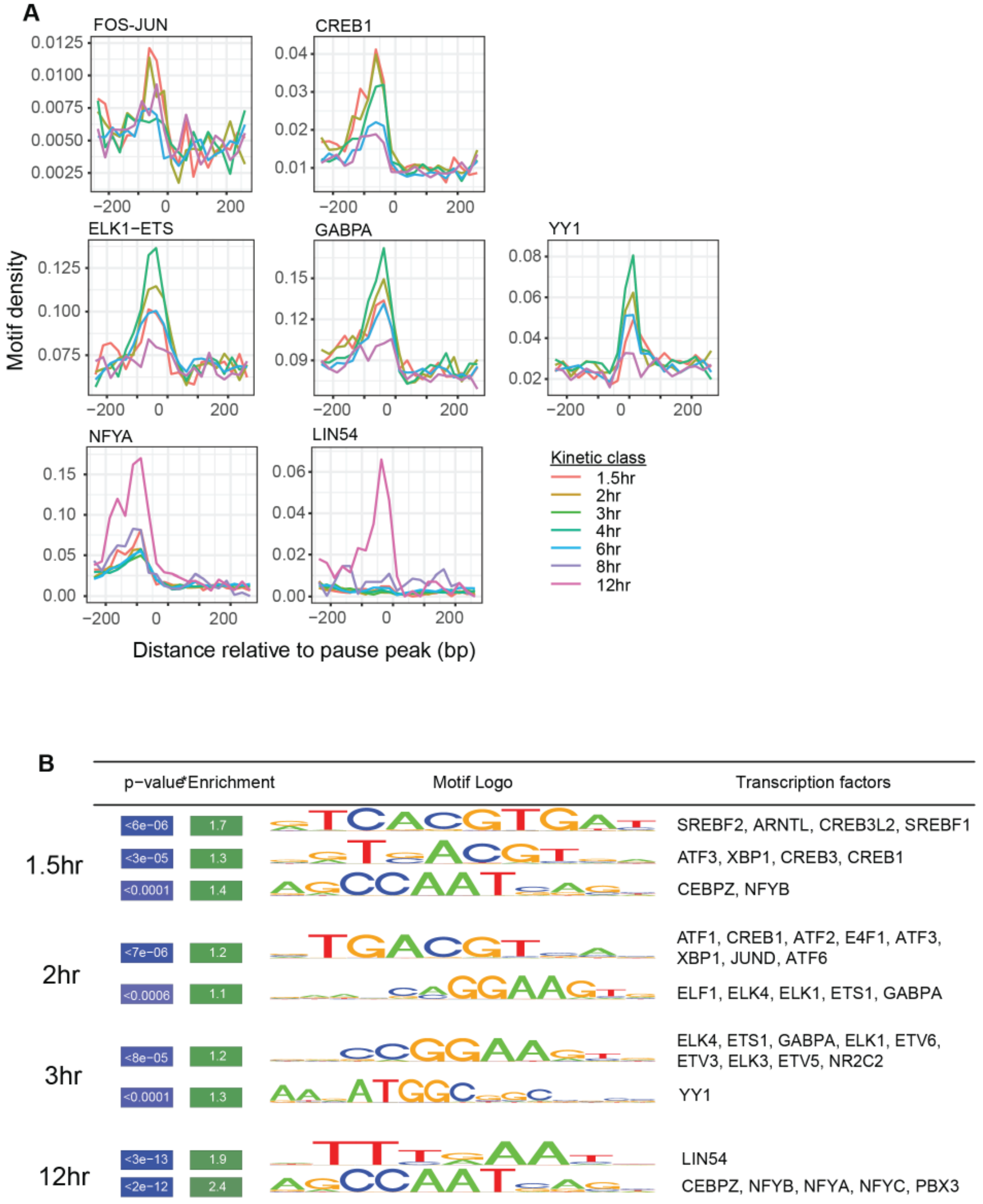


**Figure S6 (related to figure 6)**

**(A)** Composite profile of sequence motifs in promoter regions of gene kinetic classes. (In some plots, 8hr and 12hr classes are omitted for clarity). **(B)** Motif enrichment analysis results in promoters of kinetic classes. Similar motifs were collapsed into the top result with all associated transcription factors listed.

**Figure S7**

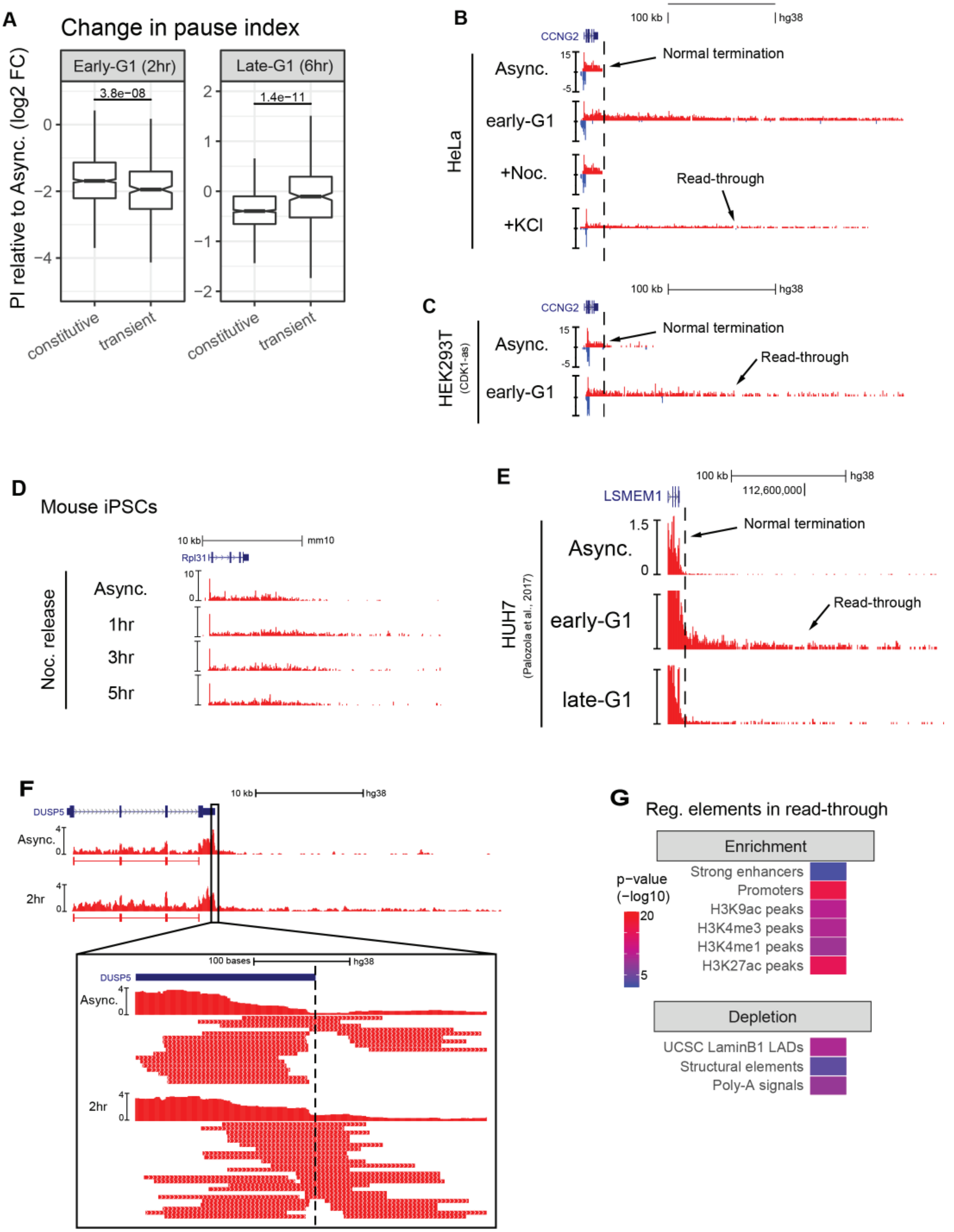
A
Change in pause index
Early-G1 (2hr)
Late-G1 (6hr)
3.8e−08
1.4e−11
PI relative to Async. (log2 FC)
constitutive
transient
B
100 kb
hg38
CCNG2
Async.
early-G1
+Noc.
+KCl
HeLa
Normal termination
Read-through
C
HEK293T
(CDK1-as)
D
Mouse iPSCs
10 kb
mm10
Rpl31
Noc. release
Async.
1hr
3hr
5hr
E
LSMEM1
112,600,000
HUH7
(Palozola et al., 2017)
late-G1
F
DUSP5
10 kb
2hr
100 bases
G
Reg. elements in read-through
Enrichment
Strong enhancers
Promoters
H3K9ac peaks
H3K4me3 peaks
H3K4me1 peaks
H3K27ac peaks
p−value
(−log10)
Depletion
UCSC LaminB1 LADs
Structural elements
Poly-A signals

**Figure S7 (related to figure 7)**

**(A)** Box plots of fold-change in pausing index relative to asynchronous cells in early gene kinetic classes during early-G1 and late-G1. **(B)** Browser tracks of a representative gene demonstrating read-through transcription during early-G1 (2hr following release from mitosis) and hyperosmotic shock (2hr treatment with 80mM KCl), but not during nocodazole treatment (2hr at 100ng/mL). Y-axis is spike-in normalized PRO-seq signal. **(C)** Browser tracks of a representative gene demonstrating read-through transcription during early-G1 (2hr following release from G2/M) in HEK293T CDK1-as cells. Y-axis is spike-in normalized PRO-seq signal. **(D)** Browser tracks of a representative gene demonstrating read-through transcription in mouse iPSCs during early-G1. Y-axis is spike-in normalized PRO-seq signal. **(E)** Browser tracks of a representative gene demonstrating read-through transcription in HUH7 cells during early-G1. **(F)** Representative browser image demonstrating full-length PRO-seq data and quantification of poly-A spanning reads in asynchronous cells and after 2hr release from mitosis. Y-axis is normalized reads per million. Below the signal track are spliced reads in BED12-format (main browser image) and individual unspliced reads in the region surrounding the annotated poly-A site (inset).

#### METHODS

##### Cell Culture

HeLa-S3 and HEK293T cell lines were cultured in Dulbecco's Modified Eagle's Medium supplemented with 10% fetal bovine serum and 1% penicillin-streptomycin at 37°C and 5% $CO_2$.

##### Construction of CDK1-AS Mutant and Cell Synchronization

A CDK1-F80A mutant was created using CRISPR/Cas9 in HEK293T cells as described in (Gravells et al., 2013). Guide oligos (5'- TGGAAAGAAACTCAAAGATGCAAA-3', 5'-CACCCATCTTTGAGTTTCTTTCCA-3') were cloned into the PX459v2 plasmid (AddGene #62988), and transfected into HEK cells with Lipofectamine 3000. To create the mutation, the following oligo donor repair template was added to the transection: 5'-AAAATGTTTGCTGGATTCTTCTCTCATATATTTTTTTTCCCCAGTCTTCAGGATGTGCTTATGCAGGATTCCAGGTTATATCTCATCGGCGAGTTTCTTTCCATGGATCTGAAGAAATACTTGGATTCTATCCCTCCTGGTCAGTACATGGATTCTTCACTTGTTAAGGTAAAAGCTTAACTAATTTTATTAATATTTAT-3'. Transfected cells were selected with 2ug/ml Puromycin for 72 hours. Surviving cells were plated in 96-well plates at 1 cell / well. Clonal lines were verified by genomic PCR and Sanger sequencing (Gravells, Tomita, et al, 2013).

##### Mitotic Synchronization and Release

HeLa-S3 cells at 25-30% confluency were treated with 2mM thymidine for 24 hours, released in fresh medium for 3 hours, then treated with 100ng/mL nocodazole for 12 hours (Whitfield, Sherlock, et al, 2002). Mitotic cells were collected by shake-off, centrifuged and washed in 1x PBS, and then either grown on 15cm dishes in fresh medium for the corresponding time or immediately permeabilized (mitotic sample). Synchronization of HEK293T CDK1-AS cells was carried out by incubation with 100uM 1NMPP-1 for 20 hours (Gravells, Tomita, et al, 2013). To release cells into mitosis, cells were carefully washed twice with pre-warmed untreated media and returned to incubation at 37C. Cells were harvested at indicated timepoints.

##### Cell Cycle Analysis

Prior to cellular permeabilization, 10% of each sample was removed and fixed in 75% cold (-20°C) ethanol. Cells were then stained with propidium iodide and DNA content was analyzed (Whitfield et al., 2002) using a BD FACS Aria II. FCS files were read into R for downstream analyses using the flowCore package. Separately, mitotic HeLa cells were also stained with DAPI and manually analyzed by microscopy in order to differentiate cells in G2 from those properly arrested in prometaphase by level of DNA condensation.

##### Cell Permeabilization

For each replicate timepoint, both floating cells in the growth medium and cells removed by scraping in 1x PBS were collected, pooled, and centrifuged at 1000xg for 5 min. Cells were resuspended in 1x PBS and 10% was removed for FACS analysis before completing the wash. 1mL of buffer P (10mM Tris-Cl pH 8.0, 10mM KCl, 250mM sucrose, 5mM $MgCl_2$, 1mM EDTA, 0.05% Tween-20, 0.5mM DTT, 10% Glycerol) was used to gently resuspend cell pellets, before adding an additional 9mL of buffer P and incubating on a shaker for 5min. Permeabilization was assessed with trypan blue and samples not initially permeabilized were again incubated on a shaker with buffer P containing 0.05% NP-40 for 5min. Permeabilized cells were then centrifuged at 1000xg for 5min before resuspension in 1mL buffer F (50mM Tris-CL pH 8.0, 40% glycerol, 5mM $MgCl_2$, 0.1 mM EDTA, 0.5mM DTT), transferred to a 1.5mL tube, and centrifuged at 1000xg for 5min. Finally, permeabilized cells were resuspended in 55μL of buffer F and 1μL of RNase-inhibitor was added before snap-freezing in liquid nitrogen and storage at -80°C.

##### PRO-seq Library Preparation

PRO-seq libraries were prepared as previously described (Mahat, Kwak, et al, 2016) with minor modifications. 0.9-4.5 x $10^6$ permeabilized cells were mixed with permeabilized *Drosophila* S2 nuclei in 4-biotin run-on reactions (1 x $10^6$ *Dm* nuclei in each A replicate and 5 x $10^4$ in each B replicate). Run-on RNA was then base-hydrolyzed for 10min on ice and enriched with an M280 streptavidin-bead binding and TRIzol extraction. Following 3'-ligation and the second bead binding, both end-repair reactions and the 5'-ligation were all performed with nascent RNA still bound to the beads (Judd et al., 2020 bioRxiv). These on-bead reactions were done in a total volume of 20μL with constant rotation before elution from the beads and the subsequent reverse transcription and PCR steps. Test amplifications were performed on 5% of the library and samples were amplified the ideal number of cycles. Following final amplification, libraries were PAGE-purified to remove adapter-dimers and select molecules below 650bp in size. Libraries were then sequenced on an Illumina NextSeq 550, producing single-end 75bp reads.

**Full-length PRO-seq Library Preparation**

Nuclei were prepared in duplicate as above from asynchronous and 2hr mitotic-release cells, totaling ~$1x10^7$ nuclei per sample. Immediately following nuclei isolation, each sample was split into two separate run-on reactions as in Mahat et al., 2016. After 5min, TRIzol was added to stop the run-on reaction and RNA was extracted. RNA was resuspended, bound to streptavidin M280 beads and extensively washed as in Mahat et al., 2016. RNA was then eluted, extracted, bound to beads again, and washed. Nascent RNA was eluted from the beads and any remaining ribosomal RNA was then removed using riboZero Gold (Illumina). Samples split at the run-on step were then pooled and 0.01ng of total Drosophila RNA (rRNA-depleted) was added as a spike-in control. Illumina TRU-seq stranded low-input RNA-seq libraries were then generated from these samples following the standard protocol and sequenced on an Illumina NextSeq 550, producing paired-end 150bp reads.

**ATAC-seq Library Preparation**

ATAC-seq was performed as previously described (Buenrostro, Giresi, et al, 2013) with minor modifications. 50,000 permeabilized cells were pelleted and then incubated for 30min in ATAC-seq reaction buffer at 37°C in a shaker at 1000rpm. DNA was then extracted with phenol-chloroform before size selection with AMPure beads. Test amplifications were performed on 5% of the library and samples were amplified the ideal number of cycles. Libraries were then sequenced on an Illumina NextSeq 550, producing single-end 75bp reads.

**PRO-seq Data Processing**

Raw fastq files were first quality trimmed (Phred score >=20) and adapter sequences removed using cutadapt (Martin, 2011). Reads were also trimmed to 36nt and those below 15nt were removed. Remaining reads were reverse complemented using the fastx-toolkit (Hannon, 2010) and then aligned with bowtie to human ribosomal sequences (GenBank: U13369.1) allowing up to 2 mismatches (Langmead, Trapnell, et al, 2009). Non-ribosomal reads were then aligned using bowtie to a combined human-drosophila (hg38 and dm6) genome allowing up to 2 mismatches while reporting only uniquely-aligning reads ("-m1" option). Alignments were then converted to a single 3'-base, which is used for all downstream data visualization and analyses. Bedgraph and BigWig files were created using BEDtools (Quinlan, and Hall, 2010) and kentUtils (Kent, Sugnet, et al, 2002), respectively. Detection of Regulatory Elements using GRO-/PRO-seq (dREG) was performed on all combined PRO-seq data to produce a list of transcription regulatory elements (TREs) (Danko et al., 2015). These regions include actively transcribed promoters and enhancers which are identified by the occurrence of bidirectional transcription.

**ATAC-seq Data Processing**

Raw fastq files were first quality trimmed (Phred score >=20) and adapter sequences removed using cutadapt. For single-end ATAC-seq data, reads were also trimmed to 36nt and those below 15nt were removed before alignment with bowtie to the human genome (hg38) allowing up to 2 mismatches while reporting only uniquely-aligning reads. Alignments were deduplicated with the picard toolkit (Picard) and extended to 200bp total length and MACS2 (Zhang, Liu, et al, 2008) was used to call broad peaks with a q-value threshold of 0.01. To reduce the effect that differences in mappability may have between libraries of different read size and single-end versus paired-end data, all analyses were limited to either our dREG regions or single-end ATAC-seq peaks, and gene annotations containing a TSS overlapping these regions. For paired-end ATAC-seq data ((Oomen, Hansen, et al, 2019): GSE121840), adapter-clipped read pairs were aligned with bowtie2 to the human genome (hg38) using a max fragment size of 2000bp. Concordant, primary alignments with high mapping quality (MAPQ>=20) were retained using samtools (Li, Handsaker, et al, 2009), deduplicated with the picard toolkit, and converted to bed files representing the full fragment for downstream visualization and analyses. Bedgraph and BigWig files were created using BEDtools and kentUtils, respectively.

**ChIP-seq Data Processing**

Raw fastq files were first quality trimmed (Phred score >=20) and adapter sequences removed using cutadapt. For single-end ChIP-seq data ((Liang, Woodfin, et al, 2015): GSE71848), reads were also trimmed to 36nt and those below 15nt were removed before alignment with bowtie to the human genome (hg38) allowing up to 2 mismatches while reporting only uniquely-aligning reads. Alignments were, deduplicated with the picard toolkit and then extended to 200bp in total for visualization and analyses. For paired-end ChIP-seq data ((Javasky, Shamir, et al, 2018): GSE108173), adapter-clipped read pairs were aligned with bowtie2 to the human genome (hg38) using a max fragment size of 800bp. Concordant, primary alignments with high mapping quality (MAPQ>=20) were retained, deduplicated with the picard toolkit, and converted to bed files representing the full fragment for downstream visualization and analyses. Bedgraph and BigWig files were created using BEDtools and kentUtils, respectively.

**GRO-seq, EU-RNA-seq, and Nascent-seq Data Processing**

Raw fastq files ((Liang, Woodfin, et al, 2015): GSE71848; (Palozola, Donahue, et al, 2017): GSE87476 (Liu, Chen, et al, 2017): GSE94479; (Kang, Shokhirev, et al, 2020): GSE141139) were first quality trimmed (Phred score >=20) and adapter sequences removed using cutadapt. Reads were also trimmed to 36nt and those below 15nt were removed. Remaining reads were aligned to human ribosomal sequences with bowtie allowing up to 2 mismatches. Non-ribosomal reads were then aligned using bowtie to the human genome (hg38). Primary alignments with high mapping quality (MAPQ>=20) were retained, deduplicated with the picard toolkit (deduplication was skipped with GRO-seq data), and converted to bed files representing the full fragment for downstream visualization and analyses. Bedgraph and BigWig files were created using BEDtools and kentUtils, respectively.

**MNase-seq Data Processing**

Raw fastq files ((Kfir, Lev-Maor, et al, 2015): GSE65644) were first trimmed to 36nt with cutadapt and then aligned with bowtie to the human genome (hg38) allowing up to 2 mismatches while reporting only uniquely-aligning reads. Alignment coordinates were converted to the predicted nucleosome center (+75bp from the start of the read) and then extended 25bp in either direction for visualization and analyses. Bedgraph and BigWig files were created using BEDtools and kentUtils, respectively.

**Full-length PRO-seq Data Processing**

Raw fastq files were first quality trimmed (Phred score >=20) and adapter sequences removed using cutadapt. Reads below 15nt were removed and remaining reads were aligned with HISAT2 (Kim, Langmead, and Salzberg, 2015) to human ribosomal sequences (GenBank:U13369.1). Non-ribosomal reads were then aligned using HISAT2 to a combined human-drosophila (hg38 and dm6) genome and low quality alignments filtered out (MAPQ >=2). Bedgraph and BigWig files were created using BEDtools and kentUtils, respectively.

#### <u>QUANTIFICATION AND STATISTICAL ANALYSIS</u>

**PRO-seq Gene Analysis**

TREs from dREG were combined with protein-coding and lincRNA annotations from GENCODE V27 for downstream analyses. Promoters were defined as TREs with any overlap by an annotated TSS and the longest corresponding gene annotation was kept. Non-promoter TREs were classified as enhancers. Polymerase pause sites were defined as the furthest upstream 50bp window containing the maximum signal within each promoter TRE. The first 500bp of annotations were excluded from gene expression analyses to avoid promoter signal and gene ends were trimmed to a maximum of 10kb to avoid potential length-bias in analyses of gene reactivation kinetics.

**PRO-seq Enhancer Analysis**

TREs that overlapped ATAC-seq peaks but did not overlap annotated protein coding or lincRNA TSSs were used as a high-confidence enhancer list. To compare enhancer activity in asynchronous and mitotic cells, these TREs were used during analyses. However, for enhancer analyses of the full timecourse, SICER (Zang, Schones, et al, 2009) was used to call transcribed regions emanating from enhancers in order to reduce the effect of lower pausing signal immediately after mitosis. SICER was run with default settings on plus and minus strand data from all timepoints pooled and then merged with enhancer TREs. Regions that overlapped either genes or read-through transcription on the same strand were removed, and the remaining regions were trimmed to a maximum 5kb. When getting counts, the TRE regions were masked in order to avoid the pause region.

**ATAC-seq Data Analysis**

NucleoATAC (Schep, Buenrostro, et al, 2015) was run on asynchronous and mitotic paired-end ATAC-seq data with defaults parameters, using peaks extended by 1kb on either side. For downstream analyses, fragments were either converted to 1bp cut-sites or binned by size; <100bp (subnucleosomal) or 180-247bp (mononucleosomal) (Buenrostro, Giresi, et al, 2013). To compare transcribed and non-transcribed regulatory elements, ATAC-seq summits were used as a reference point in order to reduce potential bias in the comparisons.

**Read-through Transcription Analysis**

Annotated protein-coding and lincRNA transcript end sites (TES's) were collected from the comprehensive Gencode GRCh38 version 27 transcripts list (159,288 transcripts in total). This list was used to select gene-ends that most closely fit our asynchronous PRO-seq data according to current models. Asynchronous data was used to compare signal in the region 5kb downstream of the TES ("termination zone") to signal further downstream in the +7.5 to +17.5kb region ("read-through region") (excluding any overlap with downstream genes). All annotations with a termination zone mappable-read density >5 RPKM and a termination zone/read-through region ratio >1 were retained. Early-G1 normalized signal (2hr timepoint) in the termination zone was also required to be at least 90% of asynchronous data to ensure only genes fully transcribed remained in the analysis. From the remaining annotations, the TES with the largest termination zone/ read-through region ratio in asynchronous data was selected (5,455 genes in total). Genes with a mappable-read density >1 RPKM in the downstream region during early-G1 were identified as having read-through transcription (3,562). To capture the full-length of the read-through, the region from the TES to the nearest downstream promoter (same-strand) was searched for the first window with a mappable-read density <0.1 RPKM.

**RNA processing analysis**

For poly-A cleavage analysis, high-confidence poly-A sites were defined based on RNA-seq data in upstream and downstream 100bp windows surrounding all annotated poly-A sites (GRC27). Poly-A sites with >1RPM in the upstream region and an upstream/downstream ratio >4 were retained. For analysis of the full-length PRO-seq data, windows were expanded to 1kb on either side of the poly-A site and poly-A spanning reads were required to cover both bases on either side of the poly-A site. Poly-A spanning reads were normalized to transcription level using the downstream signal. For splicing analysis, counts of both spliced and unspliced full-length PRO-seq reads were obtained in gene body and read-through regions. Counts were then normalized to reads per million (RPM).

**Region enrichment analysis**

Enrichment of genomic elements within regions was calculated using Locus Overlap analysis (LOLA) (Sheffield, and Bock, 2016). General classes of promoters, enhancers, poised enhancers, and CTCF-sites were created using a combination of annotated TSSs, dREG, and H3K4me1/CTCF ChIP-seq (Javasky, Shamir, et al, 2018). Enrichment analysis was performed using these groups as background with corresponding subsets of each group. For enrichment within read-through regions, genes without an early-G1 extension were expanded to the nearest downstream promoter (maximum of 50kb) and included in the background regions. Genomic elements demonstrating enrichment (or depletion) with adjusted p-values < 0.001 were included in the resulting plots.

**Gene ontology analysis**

The 'clusterProfiler' package (Yu, Wang, et al, 2012) was used in R to calculate GO-term enrichment scores for gene groups, using all active genes as background. The 'rGREAT' package (McLean, Bristor, et al, 2010) was used to assign enhancers to nearby genes and perform GO analysis in 'basal plus extension' mode. Enriched GO terms with adjusted p-values < 0.001 were retained and GO terms were then grouped by keywords (Supplemental Table S1).

**Heatmaps and composite profiles**

Count matrices for heatmaps and composite profiles were generated using deepTools (Ramirez, Ryan, et al, 2016). Data in bigwig format was binned using the computeMatrix command with 10bp bins for plotting promoter regions and 500bp bins when plotting both gene and read-through regions.

**Differential Expression Analysis to Identify Mitotic-Enriched Genes**

To compare a variety of nascent transcription datasets across multiple cell types in mitosis we pre-processed and aligned all data in the same manner. A standard differential gene expression analysis was performed using EdgeR (Robinson, McCarthy, and Smyth, 2010) to identify mitotic-enriched genes in each cell type relative to asynchronous expression. For each active gene in a given cell type, the isoform with highest magnitude fold change was retained and the resulting tables were merged across all cell types. Mitotic-enriched genes were identified by having a log2 fold-change >1.5 relative to asynchronous cells and an adjusted p-value <0.05.

**Motif Analyses**

Motif composite profiles were created by downloading BED files of motif locations throughout hg38 from PWMTools (http://ccg.vital-it.ch/pwmtools). Regions of a fixed-size are then windowed using a 1bp window with BEDtools and counts of promoter overlap are obtained in each window. Windows are summed across all regions and then divided by the total number of regions to get motif frequency. The TATA box was detected in promoters of mitotic-active genes using MEME suite tools (Bailey, Boden, et al, 2009). Promoter motif enrichment in gene kinetic classes was calculated using the rtfbsdb package using all active promoters as a background list and after correcting for GC-content. TF motifs enriched with a p-value <0.01 were retained and similar motifs were collapsed to the most significant member with individual TFs listed.

**Statistical Methods**

Comparisons of all median values were performed using the two-sided Wilcoxson's rank sum test. K-means clustering was performed in R with default settings for grouping early genes using spike-in normalized signal relative to max in the timepoints after initial gene reactivation. A two-sided Fisher's exact test was used to calculate significance of enrichment between gene and enhancer assignments by kinetic class as well as enrichment of ChIP-seq peaks in genes (promoters) and enhancers by kinetic class.
